# Disentangling group specific QTL allele effects from genetic background epistasis using admixed individuals in GWAS: an application to maize flowering

**DOI:** 10.1101/669721

**Authors:** Rio Simon, Mary-Huard Tristan, Moreau Laurence, Bauland Cyril, Palaffre Carine, Madur Delphine, Combes Valérie, Charcosset Alain

**Affiliations:** GQE - Le Moulon, INRA, Univ. Paris-Sud, CNRS, AgroParisTech, Université Paris-Saclay, 91190, Gif-sur-Yvette, France; MIA, INRA, AgroParisTech, Université Paris-Saclay, 75005, Paris, France; UE 0394 SMH, INRA, 2297 Route de l’INRA, 40390, Saint-Martin-de-Hinx, France

## Abstract

When handling a structured population in association mapping, group-specific allele effects may be observed at quantitative trait loci (QTLs) for several reasons: (i) a different linkage disequilibrium (LD) between SNPs and QTLs across groups, (ii) group-specific genetic mutations in QTL regions, and/or (iii) epistatic interactions between QTLs and other loci that have differentiated allele frequencies between groups. We present here a new genome-wide association (GWAS) approach to identify QTLs exhibiting such group-specific allele effects. We developed genetic materials including admixed progeny from different genetic groups with known genome-wide ancestries (local admixture). A dedicated statistical methodology was developed to analyze pure and admixed individuals jointly, allowing one to disentangle the factors causing the heterogeneity of allele effects across groups. This approach was applied to maize by developing an inbred “Flint-Dent” panel including admixed individuals that was evaluated for flowering time. Several associations were detected revealing a wide range of configurations of allele effects, both at known flowering QTLs (*Vgt1*, *Vgt2* and *Vgt3*) and new loci. We found several QTLs whose effect depended on the group ancestry of alleles while others interacted with the genetic background. The existence of directional epistasis was highlighted by comparing admixed with pure individuals and was consistent with epistatic interactions identified at the level of QTLs. Our GWAS approach provides useful information on the stability of QTL effects across genetic groups and can be applied to a wide range of species.

**Author summary:** Identification of genomic regions involved in genetic architecture of traits has become commonplace in quantitative genetics studies. Genetic structure is a common feature in human, animal and plant species and most current methods target genomic regions whose effects on traits are conserved between genetic groups. However, a heterogeneity of allele effects may be observed due to different factors: a group-specific correlation between the alleles of the tagged marker and those of the causal variant, a group-specific mutation at the causal variant or an epistatic interaction between the causal variant and the genetic background. We propose a new method adapted to structured populations including admixed individuals, which aims to identify these genomic regions and to unravel the previous factors. method was applied to a maize inbred diversity panel including lines from the dent and the flint genetic groups, as well as admixed lines, evaluated for flowering time. Several genomic regions were detected with various configurations of allele effects, with evidence of epistatic interactions between some of the loci and the genetic background.

## Introduction

Quantitative traits are genetically determined by numerous regions of the genome, also known as quantitative trait loci (QTLs). The advent of high density genotyping of single nucleotide polymorphism (SNPs) has opened the way to the identification of QTLs in diversity panels. These studies, referred to as genome-wide association studies (GWAS), use the linkage disequilibrium (LD) between the SNPs and the QTLs underlying the traits of interest. The panels evaluated in GWAS often include sets of individuals with complex pedigrees or genetic structure [1]. The latter is a common feature in human, animal and plant species and arises when groups of individuals cease to mate with each other and start to be subjected to different evolutionary forces.

Applying GWAS in a diversity panel including individuals from different groups raises the issue of spurious associations. The stratification of a population into genetic groups generates LD between loci that are differentiated between groups but not necessarily genetically linked. When a given trait is characterized by contrasted group-specific means, all these SNPs will correlate to it and may be detected as false positives. An efficient control of these spurious associations can be done by taking structure and kinship into account in the statistical model [1, 2]. This procedure will however limit the statistical power at differentiated SNPs, making them difficult to detect in multi-group GWAS, especially in case of rare alleles [3].

In a structured population, group-specific allele effects can be observed at SNPs, and testing an overall effect using a standard GWAS model may not be effective if the QTL effect is of opposite sign in the different groups. Such effects can result from group differences in LD between SNPs and QTLs across genetic groups. A different LD extent or linkage phase between linked loci can be explained by specific dynamics of population size such as bottlenecks or expansions [4, 5]. Such patterns of LD were identified in numerous species including human [6, 7], dairy and beef cattle [8, 9], pig [10], wheat [11] and maize [12–15]. A genetic mutation appearing in a QTL region may also lead to group-specific allele effects if it occurred in a founder specific of the genetic group. Several Mendelian syndromes of obesity where shown to result from mutation within specific ethnicities in human [16]. Another possibility consists in QTLs interacting with other loci that have differentiated allele frequencies between groups (i.e. interacting with the genetic background). In human, this possibility was discussed for a candidate gene associated with a higher risk of myocardial infarction in African American than in European populations [17, 18]. Another example is a SNP in the promoter region of *HNF4A* gene which was associated with a higher risk of developing type 2 diabetes in Askenazi compared to United Kingdom populations [19]. This locus was later proven to be interacting with another gene in the Askenazi population [20]. In maize, evidences of QTLs with group-specific allele effects can also be found, even though the cause of these differences remains unclear. The presence of allelic series has been demonstrated for QTLs associated with flowering time, including *Vgt1* [21]. A QTL with group-specific allele effects was also identified in a maize diversity panel for a phenology trait [22]. More generally, studying the stability of QTL allele effects across genetic backgrounds is an important issue. In human, it determines the ability of a genetic marker to predict the predisposition of an individual to develop a genetic disease across ethnic groups. In plant or animal breeding, it conditions the success of introgressing a favorable allele coming from a source of diversity into an elite genetic material.

Different GWAS strategies were adopted to address this issue depending on the species. In human, GWAS mostly focused on a specific genetic group, and these group-specific studies were compared later through meta-analyses [23, 24]. Some of these meta-analyses revealed highly conserved effects between populations [25, 26] while other put in evidence more differences [27]. In dairy cattle, the first GWAS studies focused on a specific breed [28–30]. More recently, multi-breed GWAS were conducted to refine QTLs locations by taking advantage of the low LD extent observed in such composite populations [31–33]. In maize, the possibility to use seeds from different origins and generations led geneticists to assemble GWAS panel with a broad range of genetic materials [34–36]. These panels often include a limited proportion of admixed individuals that were derived from crosses between individuals from different genetic groups. The genomes of these admixed individuals consist in mosaics of fragments with different ancestries. Admixture events are a common feature in living species and can contribute to the successful colonization of new environments [37, 38]. In plants, innovative admixed genetic materials were created to enable high statistical power of QTL detection along with a wide spectrum of genetic diversity studied, such as nested association mapping (NAM) [39] or multi-parent advanced generation inter-cross (MAGIC) [40]. Both NAM and MAGIC populations are of great interest to study the stability of QTL effects in a wide range of genetic backgrounds. However, they generally include a limited number of founders and do not address the stability of QTL allele effects across genetic groups.

This study aimed at evaluating the interest of producing admixed individuals, derived from a large set of parents, in order to decipher the genetic architecture of a trait using innovative GWAS models. The objectives were (i) to demonstrate the interest of multi-group analyses to identify new QTLs, (ii) to highlight the interest of applying multi-group GWAS models to identify group-specific allele effects at QTLs and (iii) to show how admixed individuals can help to disentangle the factors causing the heterogeneity of allele effects across groups: local genomic differences or epistatic interactions between QTLs and the genetic background. To our knowledge, no method has been proposed in the literature to address the last objective. This method was applied to a maize inbred population evaluated for flowering traits, including dent, flint and admixed lines. Maize flowering time is an interesting trait to analyze in quantitative genetics studies. It is considered as a major adaptive trait by tailoring vegetative and reproductive growth phases to local environmental conditions. Admixed individuals were also used to investigate the existence of directional epistasis using a test based on the mean of admixed individuals relative to that of their progenitors.

## Materials and methods

### Genetic material and genotypic data

Genetic material consisted in a panel of 970 maize inbred lines assembled within the “Amaizing” project. It gathered 300 dent lines, 304 flint lines and 366 admixed doubled haploids, further referred to as admixed lines. The dent lines were those included in the “Amaizing Dent” panel [41] and the flint lines were those included in the “CF-Flint” panel [15]. The dent and flint lines aimed at representing the diversity of their respective heterotic group used in European breeding and included several breeding generations. The admixed lines were derived from 206 hybrids between flint and dent lines, mated according to a sparse factorial design (Fig 1), followed by in situ gynogenesis [42] to produce fixed admixed inbred lines. Each dent or flint line was involved in 0 to 11 hybrids (1.21 in average), each leading to 1 to 4 admixed lines (1.77 in average). In total, 171 dent lines and 172 flint lines were involved as parents of admixed lines.

**Fig 1.**
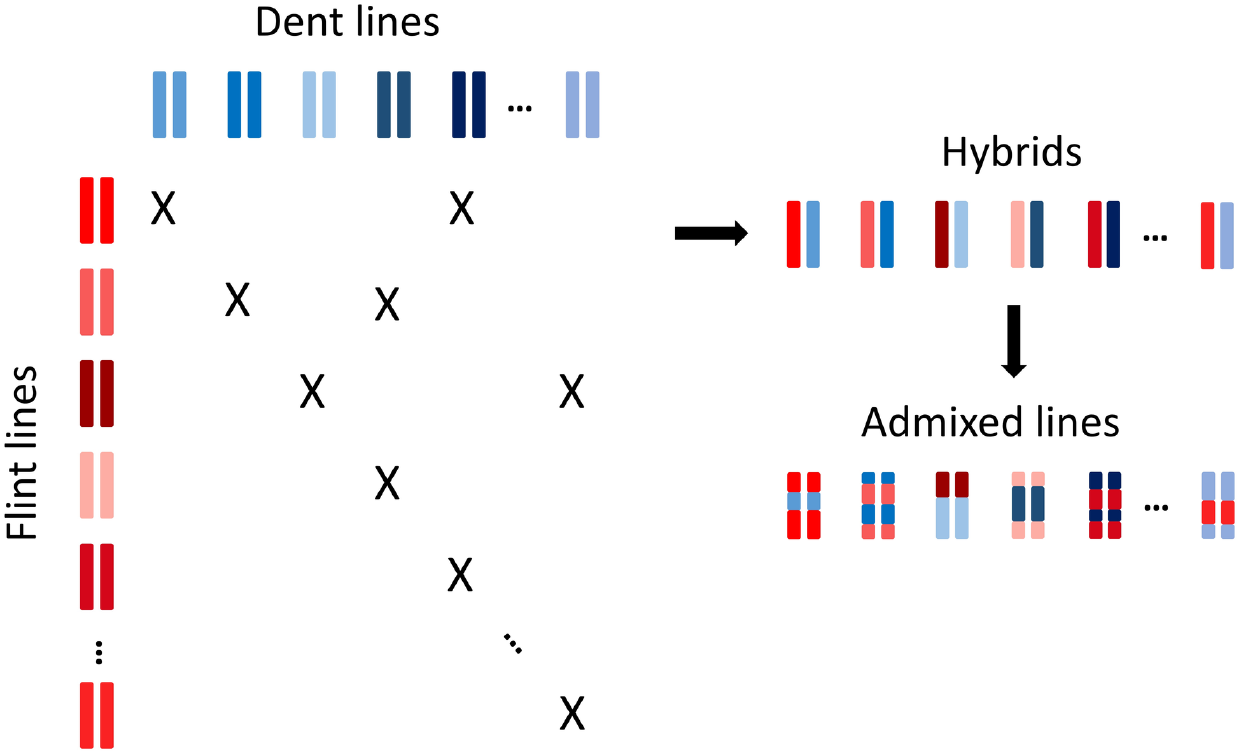
Diagram of admixed lines production from hybrids obtained by mating dent and flint lines according to a sparse factorial design

All the flint and dent lines were genotyped using the 600K Affymetrix Maize Genotyping Array [43]. Residual heterozygous data was treated as missing and all missing values were imputed independently within each group using Beagle v.3.3.2 and default parameters (Browning and Browning 2009). The admixed lines were genotyped with a 15K chip provided by the private company Limagrain which included a reduced set of SNPs from the 50K Illumina MaizeSNP50 BeadChip [44]. Eight check lines were included in both datasets to standardize the allele coding (0/1) on the common SNPs (around 9,000). The following procedure to impute admixed genotypes up to 600K SNPs is illustrated in S1 Fig. The positions of recombination breakpoints and the parental origin of the alleles for admixed lines were determined with these common SNPs. A smoothing of parental allele origins was performed for the few SNPs indicating discordant information with respect to the chromosome block in which they were located. In this case, we considered the underlying genotypic datapoint as missing. Parental origins of alleles in admixed lines were imputed up to 600K using adjacent SNP information. If a set of SNPs to be imputed was located within a recombination interval, the new position of the breakpoint was positioned at half of that ordered set, according to the physical position of the SNPs along the chromosome. Alleles at SNPs were then imputed based on their origin using parental genotypic data. The MITE associated with the flowering QTL *Vgt1* [45, 46] was also genotyped for all the individuals (0: absence, 1: presence). There was a total of 482,013 polymorphic SNPs in this dataset, for which we had information for each individual concerning the SNP allele (0/1), its ancestry (dent/flint) and the genetic background (dent/flint/admixed) in which it was observed.

The dent genome proportion of the admixed lines ranged from 0.16 to 0.86 with a mean equal to 0.51 (S2 Fig). Possible selection biases were studied along the genome by comparing the observed allele frequencies with the expected allele frequencies given the pedigree. No major pattern was observed, suggesting no or minor selection biases among the admixed lines (S3 Fig). A PCoA was performed on genetic distances computed as *D*_*i*,*j*_ = 1 − *K*_*i*,*j*_, with *K*_*i*,*j*_ being the kinship coefficient between lines *i* and *j* computed following Eq (2, see below). The flint and dent lines are clearly distinguished on the two principal components, with a small overlapping region in the center of the graph, while the admixed lines fill the genetic space between the two groups (Fig 2).

**Fig 2.**
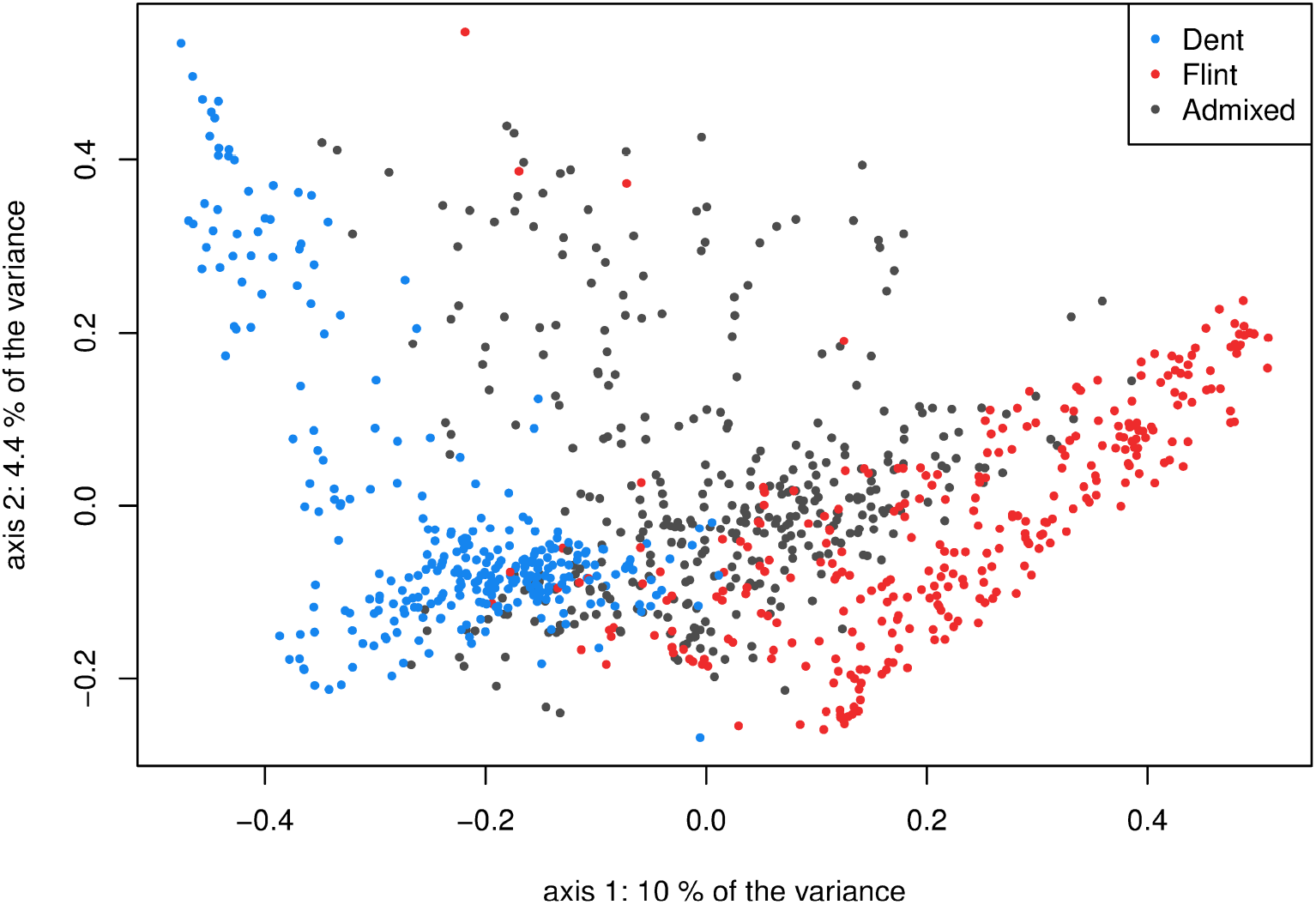
PCoA on genetic distances with coloration of individuals depending on their type: dent, flint or admixed lines

LD between pairs of loci was estimated separately in the dent and the flint datasets using two estimators. The first was the standard measure of LD, computed as the square correlation between pairs of loci *r*^2^. The second was the estimator 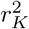 proposed by [47], accounting for relatedness estimated using Eq (2). We only considered SNPs for which at least ten individuals carried the minor allele in both dent and flint datasets. LD extent was compared between groups using both estimators. A sliding window of 1Kbp up to 2 Mbp was used to group pairs of loci with similar physical distances. The average LD was computed within each group-specific windows and revealed a higher LD extent in the dent than in the flint genetic group (S4 Fig), which was consistent with previous studies [12–15]. As suggested by [8], the persistence of LD linkage phases across flint and dent genetic groups was evaluated by computing the correlation between the *r* (and *r*_*K*_) estimated in each group using a sliding window of 1Kbp up to 2 Mbp. We also studied the consistency of marker phases between groups by computing, for each LD estimator, the correlation between their signs in the two groups. LD phases were very consistent over short physical distances but began to diverge dramatically when the loci were distant by more than 100-200 Kbp (S5 Fig).

### Phenotypic data

All the lines were evaluated *per se* at Saint-Martin-de-Hinx (France) in 2015 and 2016 for male flowering (MF) and female flowering (FF), in calendar days after sowing. Each plot consisted in a row of 25 plants. MF and FF were measured as a median value within the whole plot. Each trial was a latinized alpha design where every line was evaluated two times on average. Field trials were divided into blocks of 36 plots each. To avoid competition between genetic backgrounds, dent, flint and admixed lines were sown in different blocks. Three check individuals were repeated in all blocks (B73, F353 and UH007).

Variance components were estimated using model:

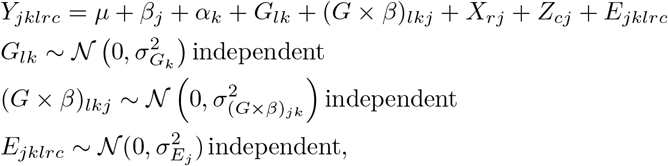

where *Y*_*jklrc*_ is the phenotype, *μ* is the intercept, *β*_*j*_ is the fixed effect of trial *j*, *α*_*k*_ is the fixed effect of genetic background *k* (dent, flint, admixed, or the different checks: B73, F353 and UH007), *G*_*lk*_ is the random genotype effect of line *l* in genetic background *k* (not for checks) with 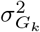 being the genotypic variance in genetic background *k*, (*G* × *β*)_*lkj*_ is the random Genotype x Environment (GxE) interaction of line *l* in genetic background *k* for trial *j*, with 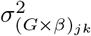 being the GxE variance in the genetic background *k* for trial *j*, *E*_*jklrc*_ is the error with 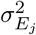 being the error variance for trial *j*, *X*_*rj*_ and *Z*_*cj*_ are the row and column random effects in trial *j* respectively, as defined by the field design. All random effects are independent of each other. The row and column effects were modeled as independent or using an autoregressive model (AR1), as determined based on the AIC criterion (S1 Table). Least squares means, further referred to as phenotypes, were computed over the whole design using the same model, with genotypes as fixed effects. Model parameters were estimated using ASReml-R and restricted maximum likelihood (ReML) [48].

### Global assessment of directional epistasis

This panel allowed us to test for the existence of directional epistasis, which refers to epistatic interactions that are biased toward high or low genetic values [49]. In the presence of directional epistatic interactions and provided no selection, we can expect the genetic mean of the admixed lines to be different from its expected value, obtained by considering only additive effects (S1 Appendix). The existence of directional epistasis was investigated using a test based on the comparison between the means of the progeny and the parental populations. The following model was applied on the joint dent, flint and admixed dataset:

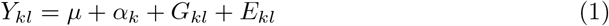

where *Y*_*kl*_ is the phenotype (least squares means) of the line among the *N* individuals of the sample, *μ* is the intercept, *α*_*k*_ is the genetic background effect with *k* ∈ {*D*, *F*, *A*} for dent, flint and admixed genetic background respectively. *G*_*kl*_ is the random genetic value of the line where ***g*** is the vector of genetic values with 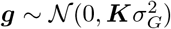, ***K*** is the kinship matrix computed following Eq (2) using allele frequencies estimated on the joint dent, flint and admixed dataset, 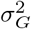 is the genetic variance, *E*_*kl*_ is the residual error of the line where ***e*** is the vector of residuals with 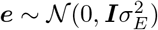, ***I*** is the identity matrix and 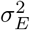 is the residual variance. For each trait, the linear combination 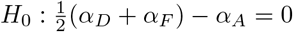 was tested to identify directional epistasis.

The kinship between individuals *i* and *j*, *K*_*ij*_, was computed following [50]:

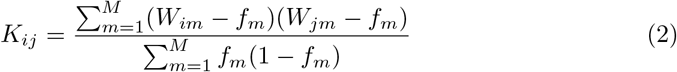

where *W*_*im*_ is the genotype of individual *i* at locus *m* coded 0/1 and *f*_*m*_ is the frequency of allele 1 at locus *m*.

### GWAS models

In this study, three GWAS models were applied to different population samples (Table 1). The GWAS strategies were (i) to analyze dent and flint lines separately using a standard GWAS model **M**_1_, (ii) to analyze dent and flint lines jointly using a GWAS model **M**_2_ accounting for allele ancestry (confounded with the genetic background) and (iii) to analyze dent, flint and admixed lines in a GWAS model **M**_3_ accounting for both allele ancestry and the genetic background of the individuals. All models aimed at detecting a SNP effect, defined as a contrast effect between alleles 0 and 1 at a given SNP.

**Table 1.** Population sample to which each GWAS model was applied with the corresponding number of SNPs conserved for the analysis (at least 10 individuals carrying the minor allelic state)

|  | Dent | Flint | Dent + Flint | Dent + Flint + Admixed |
| --- | --- | --- | --- | --- |
| $\mathbf{M}_1$ | ✓ (247,759) | ✓ (282,278) | ✗ | ✗ |
| $\mathbf{M}_2$ | - | - | ✓ (288,093) | ✗ |
| $\mathbf{M}_3$ | - | - | - | ✓(256,951) |

✓ : model was applied to the sample
✗ : model was not applied to the sample but can theoretically be, provided the addition of a genetic background effect
- : model cannot be applied to the sample or would simplify into another model

Note that the number of SNP in multi-group GWAS (**M**_2_, **M**_3_) is higher than the minimum number of SNPs in single group GWAS (**M**_1_ (Dent)). SNPs carrying redundant information within a single group were indeed reduced to a single SNP for M1 and may no longer carry redundant information when datasets are pooled (**M**_2_, **M**_3_)

### Standard GWAS model **M**_1_

The first GWAS model **M**_1_ [1] was applied separately to the dent and flint datasets. For each SNP among the *M* loci, one has:

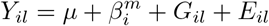

where 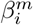 is the effect of the SNP allele *i* at locus *m* (Table 2). All other terms are identical to those described Eq (1), and the kinship was computed following Eq (2) using allele frequencies estimated for each dataset. The existence of a SNP effect was tested using hypothesis 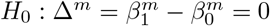.

**Table 2.** Allelic states observed in each GWAS model, resulting from a combination of SNP alleles, their ancestry and the genetic background in which they are observed.

|  | SNP | Ancestry | Genetic background | Allelic states |
| --- | --- | --- | --- | --- |
| **M<sub>1</sub>** | {0, 1} | - | - | {0, 1} |
| **M<sub>2</sub>** | {0, 1} | {D, F}<sup>a</sup> | - | {0D, 1D, 0F, 1F} |
| **M<sub>3</sub>** | {0, 1} | {D, F} | {D, A, F} | {0DD, 1DD, 0DA, 1DA, 0FA, 1FA, 0FF, 1FF} |

0 : SNP reference allele
1 : SNP alternative allele
D : Dent ancestry or genetic background
F : Flint ancestry or genetic background
A : Admixed genetic background

### Multi-group GWAS model M_2_

We applied a multi-group GWAS model **M**_2_ jointly to the flint and dent datasets, specifying the allele ancestry (confounded with the genetic background). For a given SNP *m*, one has:

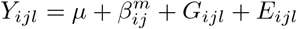

where 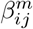 is the effect of the SNP allele *i* with ancestry *j* at locus *m*, as defined in Table 2. All other terms are identical to those described Eq. (1), and the kinship was computed following Eq (2) using allele frequencies estimated on the joint dent and flint dataset. At a given SNP, the following hypotheses were tested:

- 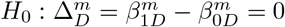
- 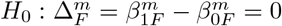
- 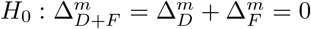
- 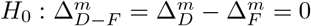

Hypotheses 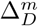 and 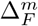 test the existence of a dent and a flint SNP effect respectively. Hypothesis 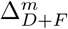 tests for a general SNP effect while 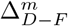 tests for a divergent SNP effect between the dent and flint ancestries

### Multi-group GWAS model M_3_

We applied a multi-group GWAS model **M**_3_ jointly to the flint, dent and admixed datasets, specifying the allele ancestry and the genetic background of the individual. For a given SNP *m*, one has:

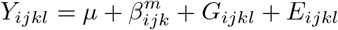

where 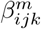 is the effect of the SNP allele *i* with ancestry *j* at locus *m* in genetic background *k*, as defined in Table 2. All other terms are identical to those described in Eq (1), and the kinship was computed following Eq (2) using allele frequencies estimated on the joint dent, flint and admixed dataset. At a given SNP, 16 hypotheses were tested (Table 3). Hypotheses referred to as “simple” (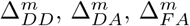 and 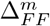) were tested to identify QTLs with a significant SNP effect for each combination of ancestries and genetic backgrounds. For instance, 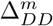 tests whether a dent SNP effect (differential effect between alleles 0 and 1 of dent ancestry) exists in the dent genetic background. Hypotheses referred to as “general” (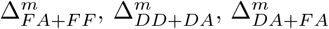, 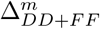 and, 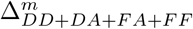) were used to identify QTLs with a mean SNP effect over ancestries and genetic backgrounds. For instance, 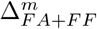 tests for a general flint SNP effect in the flint and the admixed genetic backgrounds and 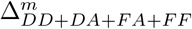 tests for a general SNP effect over ancestries and genetic backgrounds. Hypotheses referred to as “divergent” (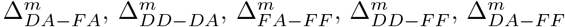, 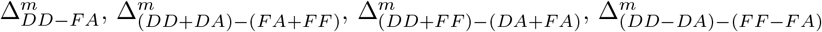) were tested to identify QTLs with a contrasted SNP effect between ancestries and/or genetic backgrounds. For instance, 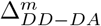 tests for a divergent dent SNP effect between the dent and the admixed genetic backgrounds, which amounts to testing an epistatic interaction between the SNP and the genetic background (see S2 Appendix for details).

**Table 3.** Linear combination tested with **M**_3_ compared to hypotheses tested using other GWAS models (**M**_1_ and **M**_2_).

| | Type | $\Delta_{DD}^m$<sup>a</sup> | $\Delta_{DA}^m$<sup>b</sup> | $\Delta_{FA}^m$<sup>c</sup> | $\Delta_{FF}^m$<sup>d</sup> | $\mathbf{M}_1$ | $\mathbf{M}_2$ |
| --- | --- | --- | --- | --- | --- | --- | --- |
| $\Delta_{DD}^m$ | simple | +1 | 0 | 0 | 0 | ✓ | ✓ |
| $\Delta_{DA}^m$ | simple | 0 | +1 | 0 | 0 | - | - |
| $\Delta_{FA}^m$ | simple | 0 | 0 | +1 | 0 | - | - |
| $\Delta_{FF}^m$ | simple | 0 | 0 | 0 | +1 | ✓ | ✓ |
| $\Delta_{FA+FF}^m$ | general | 0 | 0 | +1 | +1 | - | - |
| $\Delta_{DD+DA}^m$ | general | +1 | +1 | 0 | 0 | - | - |
| $\Delta_{DA+FA}^m$ | general | 0 | +1 | +1 | 0 | - | - |
| $\Delta_{DD+FF}^m$ | general | +1 | 0 | 0 | +1 | - | ✓ |
| $\Delta_{DD+DA+FA+FF}^m$ | general | +1 | +1 | +1 | +1 | - | - |
| $\Delta_{DA-FA}^m$ | divergent | 0 | +1 | -1 | 0 | - | - |
| $\Delta_{DD-DA}^m$ | divergent | +1 | -1 | 0 | 0 | - | - |
| $\Delta_{FA-FF}^m$ | divergent | 0 | 0 | +1 | -1 | - | - |
| $\Delta_{DD-FF}^m$ | divergent | +1 | 0 | 0 | -1 | - | ✓ |
| $\Delta_{(DD+DA)-(FA+FF)}^m$ | divergent | +1 | +1 | -1 | -1 | - | - |
| $\Delta_{(DD+FF)-(DA+FA)}^m$ | divergent | +1 | -1 | -1 | +1 | - | - |
| $\Delta_{(DD-DA)-(FF-FA)}^m$ | divergent | +1 | -1 | +1 | -1 | - | - |

<sup>a</sup> $\Delta_{DD}^m = \beta_{1DD}^m - \beta_{0DD}^m$
<sup>b</sup> $\Delta_{DA}^m = \beta_{1DA}^m - \beta_{0DA}^m$
<sup>c</sup> $\Delta_{FA}^m = \beta_{1FA}^m - \beta_{0FA}^m$
<sup>d</sup> $\Delta_{FF}^m = \beta_{1FF}^m - \beta_{0FF}^m$
✓ : hypothesis also tested using the corresponding GWAS model
- : hypothesis not tested using the corresponding GWAS model

On a biological standpoint, a QTL with contrasted SNP effects between groups can be caused by (i) a local genomic difference due to a group-specific genetic mutation and/or to group differences in LD or (ii) an interaction with the genetic background. Under the first hypothesis, one expects that the effect of a SNP depends on its ancestry but not on the genetic background (admixed or pure, see Fig 3-**a**). Under the second hypothesis, we expect a SNP effect, for a given ancestry, to vary depending on the genetic background. One example would be a QTL with a strong SNP effect in a dent genetic background, but none in the flint genetic background, while the SNP effects would be of intermediate size for alleles of both ancestries in the admixed genetic background (see Fig 3-**b**). Note that other complex configurations are possible, justifying the inclusion of all tests in the analysis.

**Fig 3.**
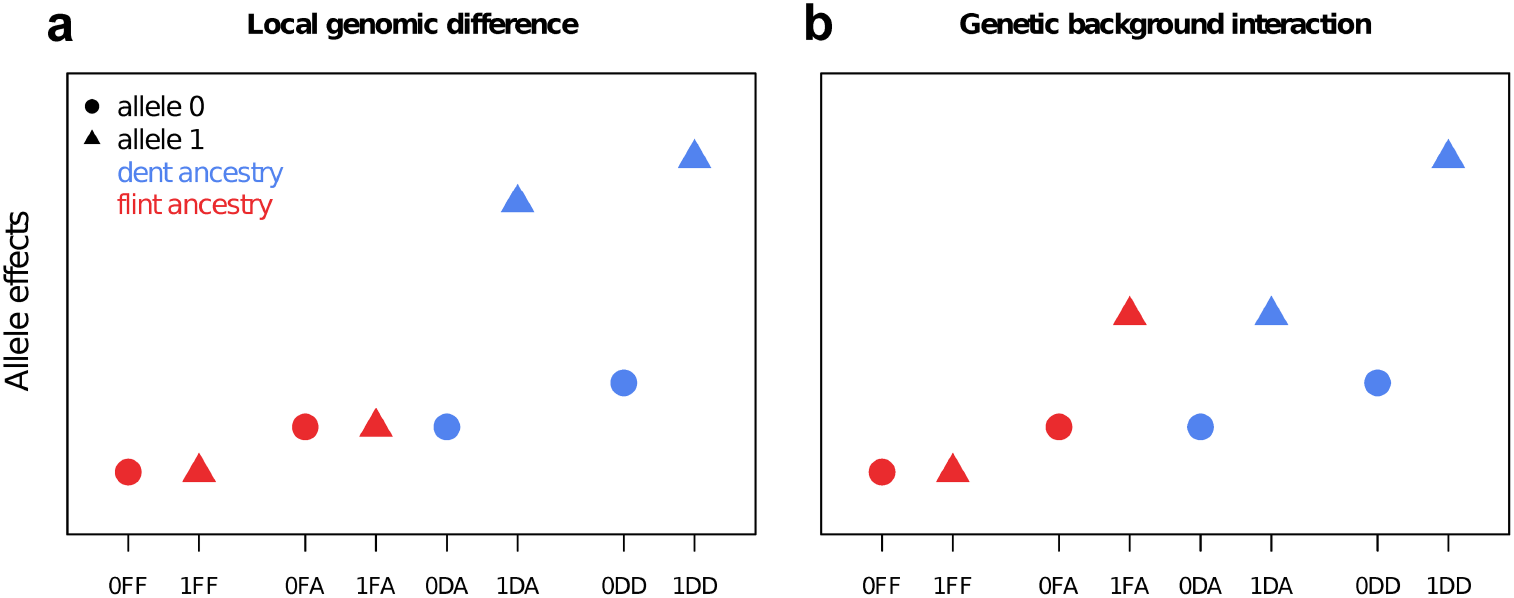
Schematic of allele effects when divergent SNP effects are observed between groups, depending on the biological hypothesis: (**a**) local genomic difference between groups and (**b**) allele effects interacting with the genetic background. The denomination of the allelic states on the x-axis include the SNP allele (0/1), its ancestry (D/F) and the genetic background in which it is observed (D/A/F), as presented in Table 2.

For the three GWAS models, a SNP was discarded if its minor allelic state (Table 2) was carried by less than 10 individuals, or if it carried a redundant information with another SNP. Model parameters were estimated using ReML and the linear combinations of fixed effects were tested using Wald tests, both implemented in the R-package MM4LMM [51]. The false discovery rate (FDR) was controlled by applying the procedure of [52] jointly to the whole set of tests defined by each GWAS strategy, and repeatedly for each trait. For a given hypothesis tested, significant SNPs were clustered into QTLs if they were located within a physical window of 3 Mbp, leading to a LD below 0.05 between markers of different QTLs.

## Results

### Phenotypic analysis and directional epistasis

We observed a substantial phenotypic variability within the dent, flint and admixed genetic backgrounds. The variance components estimated in the phenotypic analysis were summarized in S1 Table. Similar trends were observed for both MF and FF. The admixed genotypic variance was lower than the dent and flint genotypic variances, which were themselves comparable. GxE variances were limited and the broad sense heritabilities were high for each genetic background, ranging from 0.88 in the admixed lines to 0.96 in the dent and flint lines for both MF and FF.

The presence of admixed lines allowed us to test the existence of directional epistasis which was significant for both MF and FF (Table. 4). The mean of admixed lines estimated using a model accounting for relatedness differed significantly from the one expected without directional epistatic interactions. On average, admixed lines flowered as late as dent lines while the flint lines flowered earlier.

**Table 4.** Test for directional epistasis with group-specific means estimated by the model (Eq.1) and the p-value (pval) of the directional epistatic deviation

|  | Dent | Flint | Admixed | pval |
| --- | --- | --- | --- | --- |
| MF | 68.26 | 66.26 | 68.44 | 3.14 $10^{-10}$ *** |
| FF | 69.84 | 67.87 | 70.16 | 2.05 $10^{-11}$ *** |

\*\*\* : $pval < 10^{-3}$ ; \*\* : $10^{-3} < pval < 10^{-2}$ ; \* : $10^{-2} < pval < 5 \times 10^{-2}$

### Associations detected and comparison of GWAS strategies

For each GWAS model, two levels of FDR were used: 5% and 20% to declare a SNP as significantly associated. The number of significant SNPs detected and the corresponding number of QTLs were summarized in Table 5 for both traits. The location of QTLs detected using a FDR of 20% was represented along the genome in Fig 4 for MF and in S6 Fig for FF. All associations were listed in S2 Table and S3 Table. Note that major QTLs detected by a model (e.g. **M**_1_) may be discarded with another model (e.g. **M**_3_) because of the filtering on allele frequencies.

**Table 5.** Number of SNPs associated with each trait, depending on the GWAS strategy, using a FDR of 5% and 20%. The number of corresponding QTLs is also indicated

|  | MF |  |  |  | FF |  |  |  |
| --- | --- | --- | --- | --- | --- | --- | --- | --- |
|  | 5% |  | 20% |  | 5% |  | 20% |  |
|  | SNP | QTL | SNP | QTL | SNP | QTL | SNP | QTL |
| $\mathbf{M}_1^a$ | **7** | **2** | **56** | **24** | **8** | **3** | **38** | **14** |
| $\Delta^m$ (Dent) | 4 | 1 | 35 | 12 | 4 | 1 | 22 | 6 |
| $\Delta^m$ (Flint) | 3 | 1 | 21 | 13 | 4 | 2 | 16 | 8 |
| $\mathbf{M}_2^a$ | **4** | **1** | **57** | **22** | **4** | **1** | **7** | **3** |
| $\Delta_D^m$ | 4 | 1 | 37 | 9 | 4 | 1 | 4 | 1 |
| $\Delta_F^m$ | - | - | 3 | 2 | - | - | 2 | 1 |
| $\Delta_{D+F}^m$ | 1 | 1 | 11 | 7 | 2 | 1 | 2 | 1 |
| $\Delta_{D-F}^m$ | - | - | 18 | 9 | - | - | 1 | 1 |
| $\mathbf{M}_3^a$ | **7** | **3** | **116** | **41** | **6** | **2** | **11** | **6** |
| $\Delta_{DD}^m$ | 4 | 1 | 32 | 8 | 4 | 1 | 4 | 1 |
| $\Delta_{DA}^m$ | - | - | 1 | 1 | - | - | - | - |
| $\Delta_{FA}^m$ | 2 | 1 | 10 | 2 | - | - | 1 | 1 |
| $\Delta_{FF}^m$ | - | - | 1 | 1 | - | - | - | - |
| $\Delta_{FA+FF}^m$ | - | - | 4 | 4 | - | - | - | - |
| $\Delta_{DA+DD}^m$ | - | - | 10 | 4 | - | - | - | - |
| $\Delta_{DA+FA}^m$ | - | - | 11 | 5 | - | - | - | - |
| $\Delta_{DD+FF}^m$ | 2 | 2 | 34 | 12 | 2 | 2 | 6 | 2 |
| $\Delta_{DD+DA+FA+FF}^m$ | - | - | 19 | 6 | 1 | 1 | 2 | 1 |
| $\Delta_{DA-FA}^m$ | - | - | 2 | 2 | - | - | - | - |
| $\Delta_{DD-DA}^m$<sup>b</sup> | - | - | 4 | 4 | - | - | - | - |
| $\Delta_{FA-FF}^m$<sup>b</sup> | - | - | 1 | 1 | - | - | - | - |
| $\Delta_{DD-FF}^m$ | - | - | 15 | 5 | - | - | - | - |
| $\Delta_{(DD+DA)-(FA+FF)}^m$ | - | - | 5 | 3 | - | - | - | - |
| $\Delta_{(DD+FF)-(DA+FA)}^m$<sup>b</sup> | - | - | 5 | 4 | - | - | 2 | 2 |
| $\Delta_{(DD-DA)-(FF-FA)}^m$<sup>b</sup> | - | - | 2 | 2 | - | - | - | - |

<sup>a</sup> number of SNPs detected over the set of tests (a given SNP can be detected using different tests)
<sup>b</sup> hypothesis testing an interaction between the QTL and the genetic background

**Fig 4.**
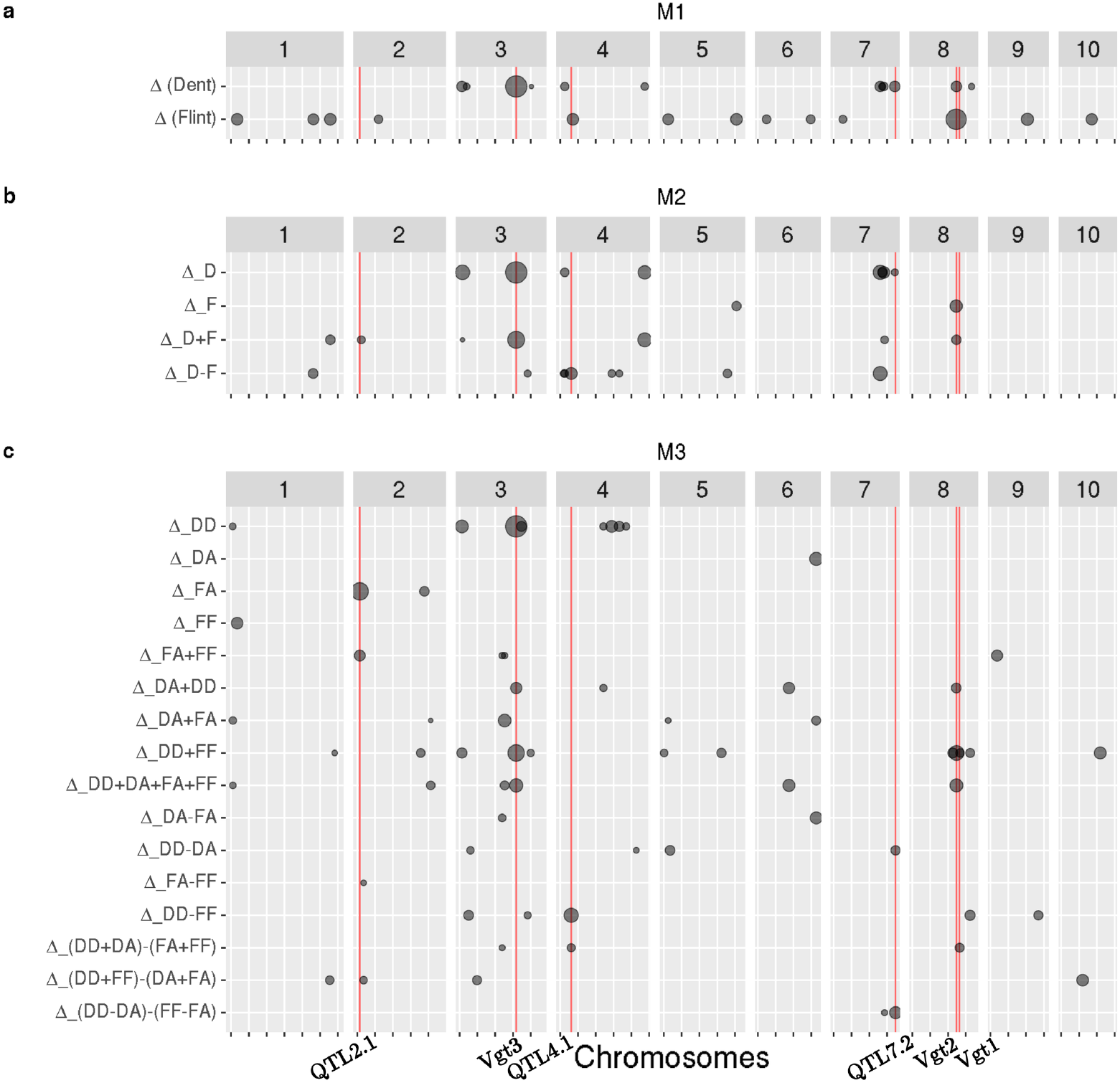
Position of QTLs detected by each GWAS strategy for MF using a FDR of 20%. The size of the grey dots is proportional to the −log_10_(pval) of the test at the most significant SNP of the region. Red vertical lines and names below correspond to QTL discussed in section “Highlighted QTLs”. Note that major QTLs detected by a model may be discarded with another model because of filtering on allele frequencies

First, a standard GWAS model **M**_1_ was applied separately to the dent and the flint datasets. Based on a 20% FDR, 35 SNPs were associated with MF in the dent dataset while 21 SNPs were associated in the flint dataset. These SNPs can be clustered into 12 QTLs in the dent dataset and into 13 QTLs in the flint dataset. Interestingly, none of these SNPs were detected in both datasets and they only pointed to one common QTL between datasets, which was located in the vicinity of Vgt2 on chromosome 8 [14].

Secondly, dent and flint datasets were analyzed jointly using model **M**_2_, which takes into account the dent or flint ancestry of the allele. Note that the allele ancestry is confounded with the genetic background in this model. Based on a 20% FDR, 57 SNPs were associated with MF and were significant for 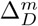 (37 SNPs), 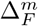 (3 SNPs), 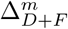 (11 SNPs) and 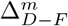 (18 SNPs). Note that some SNPs displayed more than one significant test, which explains why the total number of SNPs over the four tests did not sum to 57. These SNPs can be clustered into 22 QTLs that were significant for 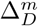 (9QTLs), 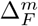 (2 QTLs), 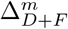 (7 QTLs) 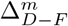 (9 QTLs). Note that some QTLs were already detected using **M**_1_ such as the QTL located in the vicinity of *Vgt3* on chromosome 3 [53, 54] detected in the dent dataset. Other QTLs were specific to **M**_1_ such as the QTL located on chromosome 2 detected in the flint dataset, or specific to **M**_2_ such as the QTL located chromosome 5 detected using 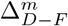. Based on a 20% FDR, a similar number of QTLs was detected between **M**_1_ and **M**_2_ for MF, while more QTLs were detected using **M**_1_ than **M**_2_ for FF.

Finally, the dent, flint and admixed lines were analyzed jointly using model **M**_3_ which distinguished the allele ancestry and the genetic background. The existence of a dent SNP effect was tested in the dent 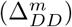 and in the admixed genetic backgrounds 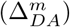, and similarly for the flint SNP effect (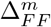 and 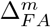). Several hypotheses on general and divergent SNP effects were also tested between ancestries and genetic backgrounds (Table 3). Based on a 20% FDR, 116 SNPs were associated with MF and were significant for 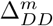 (32 SNPs), 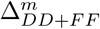 (34 SNPs), 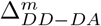 (4 SNPs), 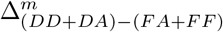 (5 SNPs) and others. These SNPs can be clustered into 41 QTLs that were significant for 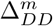 (8 QTLs), 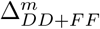 (12 QTLs), 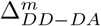 (4 QTLs), 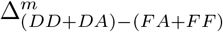 (3 QTLs) and others. Note that some of the QTLs were already detected using **M**_1_ and **M**_2_ such as the QTL located in the vicinity of *Vgt3* on chromosome 3, while several QTLs were specific to **M**_3_ such as the two QTLs detected in chromosome 2 using 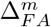. Several QTLs were detected as showing a divergent SNP effect, including hypotheses testing an interaction with the genetic background. Based on 5% and 20% FDRs, the number of QTLs detected with **M**_3_ was the highest for MF and intermediate between **M**_1_ and **M**_2_ for FF.

### Highlighted QTLs

Among the 41 QTLs detected for MF with **M**_3_, six QTLs were selected and studied in further details. The five first QTLs had (i) at least one significant test among **M**_3_ hypotheses based on a FDR of 20%, and (ii) a large frequency for each allele with a minimum of 30 lines carrying the minor allelic state (*QTL7.2*). Among them, one SNP was located in the vicinity of *Vgt2* [14] and another in the vicinity of *Vgt3* [53, 54]. In addition to these five QTLs, we also considered a MITE polymorphism known to be associated with *Vgt1*, a flowering QTL detected in several studies [21, 45, 46]. For all QTLs, information concerning their physical position along the genome, the frequency of each allelic state and their −log_10_(pval) at each test was summarized in Table 6. The distribution of the phenotypes is illustrated for each allele after adjusting the variation due to the polygenic background in Fig. 5, and their location along the genome is indicated by red vertical lines in Fig. 4. Other QTLs had interesting profiles, showing either group-specific allele effects conserved between ancestries or interactions with the genetic background, and are presented in S7 Fig and S4 Table.

**Table 6.**
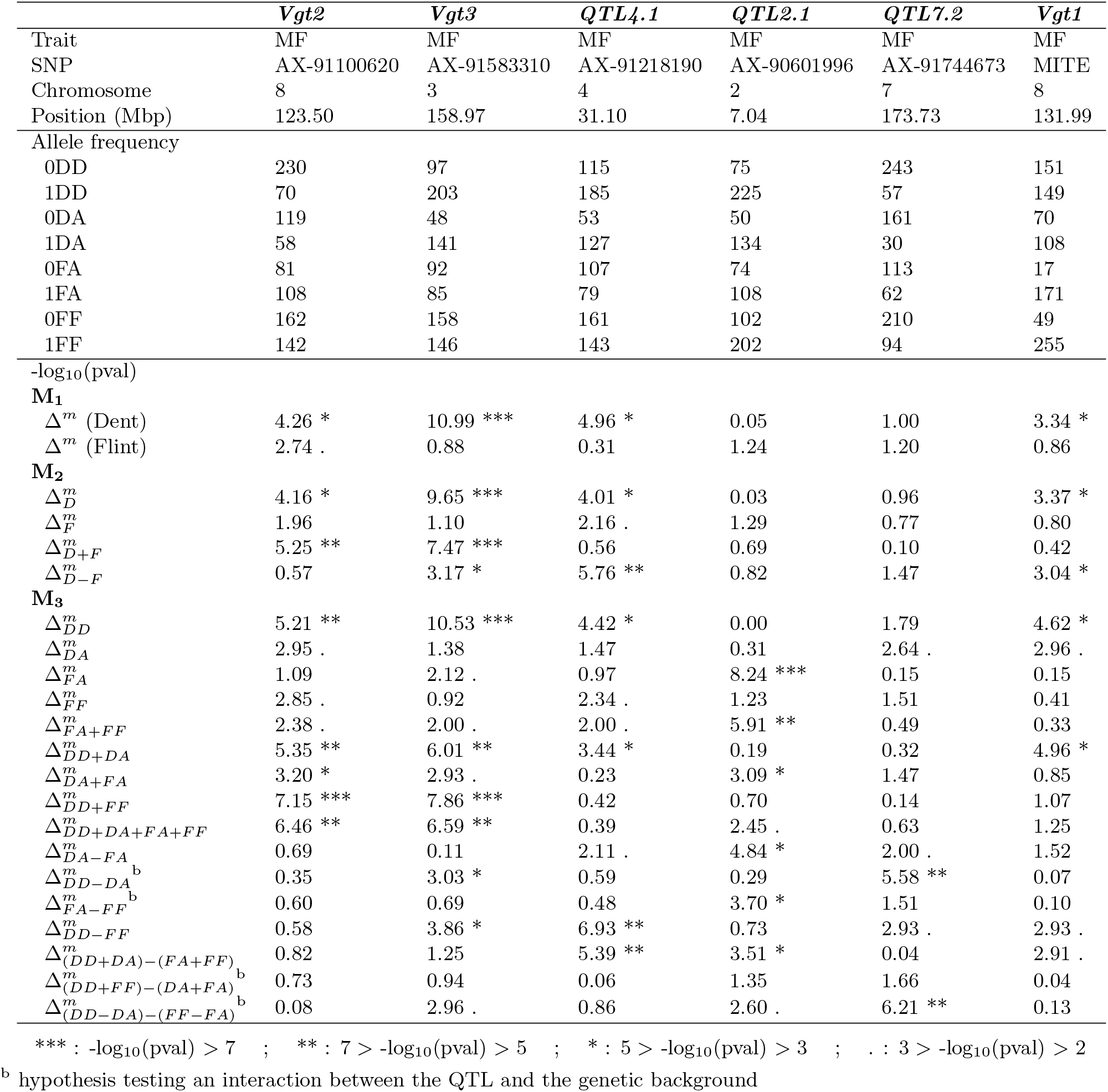
Information regarding the six highlighted QTLs

**Fig 5.**
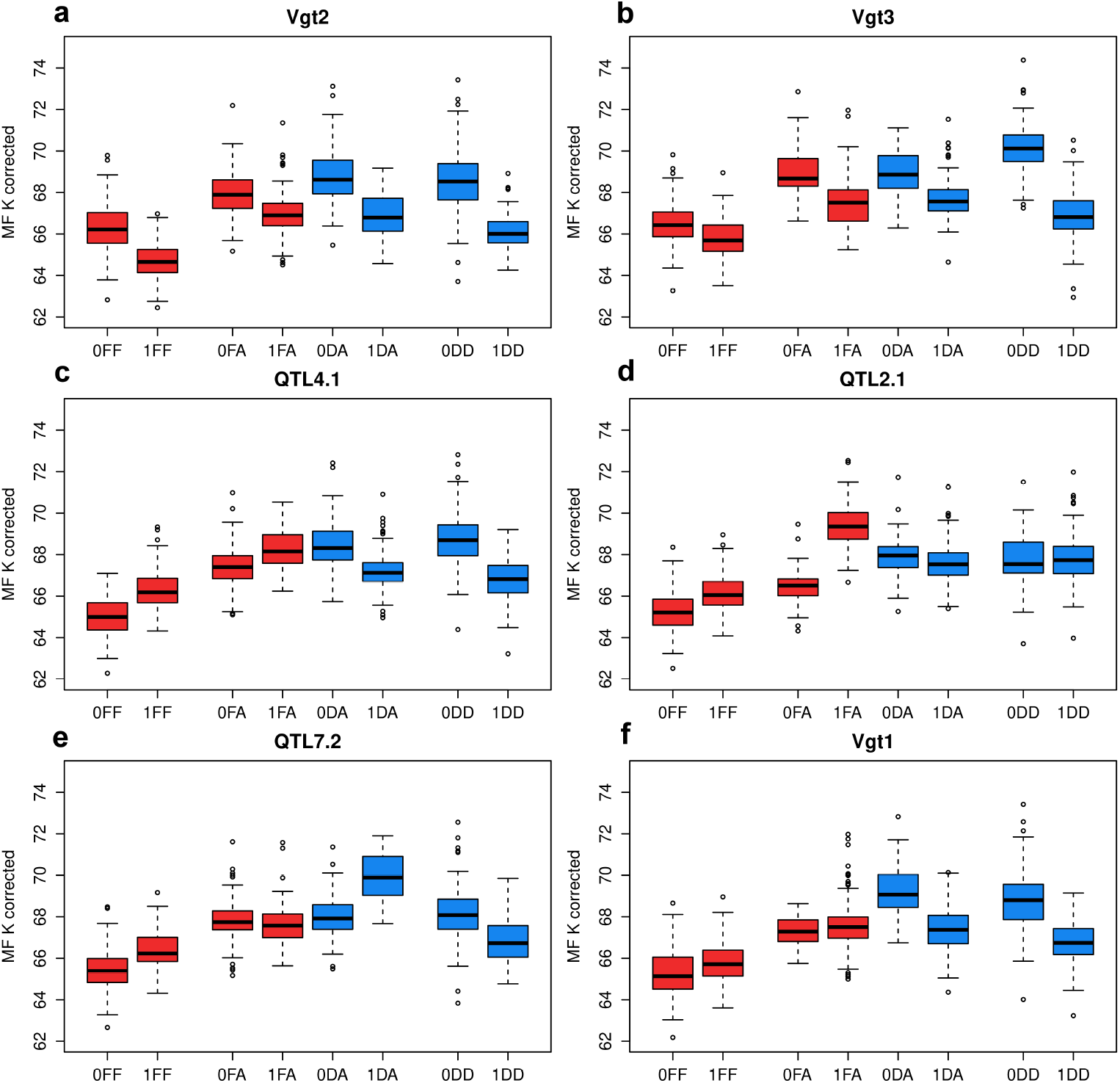
Boxplots of phenotypes adjusted for polygenic background variation using relatedness (MF K corrected) for the different alleles of the six highlighted QTLs: (**a**) *Vgt2*, (**b**) *Vgt3*, (**c**) *QTL4.1*, (**d**) *QTL2.1*, (**e**) *QTL7.2* and (**f**) *Vgt1* using **M**_3_. The denomination of the allelic states on the x-axis includes the SNP allele (0/1), its ancestry (D/F) and the genetic background in which it was observed (D/A/F), as presented in Table 2

The SNP matching *Vgt2* region on chromosome 8 was detected as associated with MF (5% FDR) using 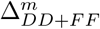 (−log_10_(pval)=7.15) in **M**_3_. This QTL showed a conserved effect across ancestries and genetic backgrounds (Fig. 5-**a**). This observation was supported by a high −log_10_(pval) for tests related to a general SNP effect: 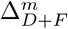 (5.25), 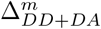 (5.35), 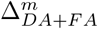 (3.20), 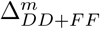 (7.15) and 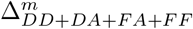 (6.46), and a low −log_10_(pval) for tests related to divergent SNP effects (all below 1).

The SNP matching *Vgt3* region on chromosome 3 was detected as associated with MF (5% FDR) using 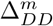 (10.53) in **M**_3_. This QTL showed a large effect in the dent genetic background, a medium effect in the admixed genetic background regardless of the allele ancestry and a small effect in the flint genetic background (Fig. 5-**b**). This observation was supported by a high −log_10_(pval) for the tests related to the dent SNP effect in the dent genetic background: Δ^*m*^ (Dent, 10.99), 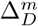 (9.65) and 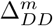 (10.53), and a low −log_10_(pval) for the tests related to the flint SNP effect in a flint genetic background. Like for *Vgt2*, a high −log_10_(pval) was also detected for tests related to a general SNP effect: 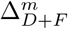 (7.47), 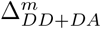 (6.01), 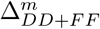 (7.86) and 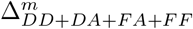 (6.59), but a high −log_10_(pval) was detected for the test related to a divergent SNP effect between the dent and the flint genetic backgrounds: 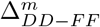 (3.86). There was also a high −log_10_(pval) for a divergent dent SNP effect between different genetic backgrounds: 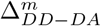 (3.03). All these results support the existence of a QTL effect that tends to be higher when the dent genome proportion increases within individuals. It suggests that *Vgt3* interacts with the genetic background for MF.

The SNP matching a region further referred to as *QTL4.1* on chromosome 4 was detected as associated with MF (20% FDR) using 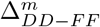 (6.93) in **M**_3_. This QTL showed a contrasted effect between alleles of different ancestries with an apparent inversion of effects (Fig. 5-**c**). This observation was supported by a high −log_10_(pval) for the tests related to a divergent SNP effect between ancestries: 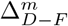 (5.76), 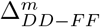 (6.93) and 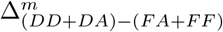 (5.39). Conversely a low −log_10_(pval) was detected for tests 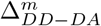 and 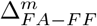, which would have otherwise suggested an interaction with the genetic background. These results support the existence of a local genomic difference at *QTL4.1* between the dent and the flint genetic groups for MF, but no interaction with the genetic background.

The SNP matching a region further referred to as *QTL2.1* on chromosome 2 was detected as associated with MF (5% FDR) using 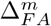 (8.24) in **M**_3_. This QTL showed a flint effect in the admixed genetic background (Fig. 5-**d**), which was supported by a high −log_10_(pval) for the test 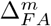 (8.24). Although there was a high −log_10_(pval) for a general flint SNP effect across genetic backgrounds: 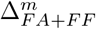 (5.91), a high −log_10_(pval) was observed for a divergent SNP effect between those same alleles: 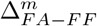 (3.70). A high −log_10_(pval) was also observed for a divergent SNP effect between different ancestries in the admixed genetic background: 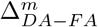 (4.84). All these results support the existence of a QTL effect existing only for alleles of flint ancestry in the admixed genetic background. It suggests that *QTL2.1* is specific of flint ancestry and interacts with the genetic background for MF.

The SNP matching a region further referred to as *QTL7.2* on chromsome 7 was detected as associated with MF (20% FDR) using 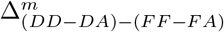 (6.21) in **M**_3_. This QTL showed contrasted dent effects between the dent and the admixed genetic backgrounds (Fig. 5-**e**). This observation was supported by a high −log_10_(pval) for the test related to a divergent dent SNP effect between genetic backgrounds: 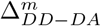 (5.58). A high −log_10_(pval) was also observed for the hypothesis testing the equality between the divergent dent SNP effect and the divergent flint SNP effect: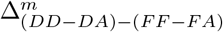 (6.21). All these results support the existence of a QTL with opposite effects between the dent and the admixed genetic backgrounds. It suggests that *QTL7.2* interacts with the genetic background for MF.

The MITE known to be associated with *Vgt1* was never detected for MF using a FDR of 5% or 20%. However, it showed a dent effect that was conserved between the dent and the admixed genetic background, and no flint effect (Fig. 5-**f**). This observation is supported by a high −log_10_(pval) for tests related to the dent SNP effect: Δ^*m*^ (Dent) (3.34), 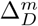 (3.37), 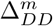 (4.62) and 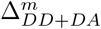 (4.96), and a low −log_10_(pval) for tests related to flint SNP effects. These results support the existence of a local genomic difference at *Vgt1* between flint and dent genetic groups but no interaction with the genetic background for MF.

## Discussion

The stratification of the population sample into distinct genetic groups is a common feature in GWAS studies. Such structure challenges the methods to detect QTLs because (i) spurious associations may be detected if the genetic structure is not accounted for by the statistical model, (ii) QTLs whose polymorphism is correlated with the genetic structure generally have a low probability of being detected when structure/relatedness is accounted for in the model, and (iii) group differences in LD, group-specific genetic mutations and/or epistatic interactions with the genetic background may prevent the detection of SNPs when testing only their average effect.

### Accounting for genetic groups in GWAS

A simple way to deal with genetic groups is to analyze them separately. In our study, a standard GWAS model **M**_1_ was applied separately within the dent and the flint datasets. High heritabilities were estimated for each genetic group in the phenotypic analysis, highlighting the suitability of these datasets to detect QTLs. Among the QTLs detected for MF, only one was detected in both dent and flint datasets, and not at the same SNPs, while none were detected in common for FF. One may question whether observing such differences between datasets indicated group specific allele effects, or simply group differences in terms of statistical power. This question often arises when GWAS is applied separately to genetic groups, as in maize [15, 55] or dairy cattle [56, 57], and is very difficult to answer except for obvious configurations such as associations at SNPs segregating only in one group.

Another way to handle genetic groups is to analyze them jointly. One possibility is to apply model **M**_1_ while specifying genetic structure as a global fixed effect, in order to prevent the detection of spurious associations. In dairy cattle, this strategy generally improved the precision concerning QTL locations by taking advantage of the low LD extent observed in multi-group datasets. However, while [33] and [32] observed a gain in statistical power due to a larger population size, [31] detected less QTLs by combining breeds compared to separate analyses. They attributed this finding to the limited amount of QTLs segregating within both Holstein and Jersey breeds, but also reported that QTLs detected in both breeds showed only small to medium correlations between within-breed estimates of SNP effects (e.g. 0.082 for milk yield). Obviously, applying **M**_1_ jointly to genetic groups does not address directly the problem of whether QTL effects are conserved or not between genetic groups.

A model specifying group specific allele effects was referred to as **M**_2_ in this study. As with **M**_1_, the existence of a dent 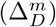 and a flint 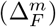 SNP effects can be tested, but **M**_2_ also allows us to test the existence of a general 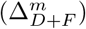 and a divergent 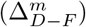 SNP effects between flint and dent ancestries. Note that testing 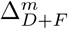 is similar, although not strictly equivalent, to testing a SNP effect by applying **M**_1_ to a multi-group dataset. Using the hypotheses specifically tested in **M**_2_ (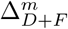 and 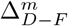), it was possible to detect new QTLs that were not detected with **M**_1_. In particular, QTLs were detected as having a divergent SNP effect between the dent and flint genetic groups, proving the existence of group-specific QTL effects in this dataset. Several QTLs were detected in common with **M**_1_ but each strategy allowed the detection of specific QTLs, demonstrating the complementarity between the models. For equivalent tests in **M**_1_ and **M**_2_ (e.g. Δ^*m*^ (Dent) in **M**_1_ and 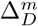 in **M**_2_), the lower number of associations detected with **M**_2_ can mostly be attributed to a more conservative filtering on allele frequencies. In conclusion, **M**_2_ was efficient to identify QTLs with either conserved or specific allele effects between ancestries, but observing group-specific allele effects provided little insight regarding the cause of this specificity. Admixed individuals can help to tackle that issue.

### Benefits from admixed individuals

Admixed individuals were generated for this study by mating pure individuals of each group according to a sparse factorial design. Integrating these admixed individuals in GWAS can be done by simply analyzing the joint multi-group dataset using **M**_1_ or **M**_2_, which may lead to a gain in statistical power, due to an increase in population size. More interestingly, admixed individuals can be used to disentangle the factors causing the heterogeneity of allele effects across groups.

We developed model **M**_3_ to distinguish the allele ancestry (dent/flint) and the genetic background (dent/flint/admixed). 41 QTLs were detected for MF (20% FDR). While many of these QTLs were previously detected using **M**_1_ and **M**_2_, the new hypotheses tested allowed us to discover new interesting regions. These new QTLs resulted from a gain in statistical power by (i) testing an overall SNP effect for SNP with conserved effects accross ancestries and/or genetic backgrounds, or by (ii) testing hypotheses for complex configurations between allele effects. The new hypotheses tested with **M**_2_ and **M**_3_ did not lead to an increase in false positive rate, based on the observation of the QQ-plots of the test p-values (results not shown).

Note that the idea of exploiting admixed individuals has been proposed in the creation of NAM [39] and MAGIC [40] populations. Compared to our approach, such experimental populations include a limited number of founders, generally selected in different genetic groups. This is beneficial to increase power of detection for alleles which were rare in the initial genetic group(s). However these populations cannot address the question of the epistatic interaction with the genetic background of the original groups, which is possible in our study thanks to the use of numerous parents. Both our approach and NAM and MAGIC designs are therefore expected to have complementary properties.

### Heterogeneity of maize flowering QTL allele effects

From a global perspective, a high number of QTLs have been detected in previous maize studies [15, 21, 36, 58, 59]. When evaluating the American and European NAMs, [21] and [60] showed that flowering time is a trait controlled by a large number of QTLs, many of which display variable effects across individual recombinant populations. Our study highlighted consistently a high number of QTLs and confirmed a large variation in effects. It provides further elements on the origin of this variation, by identifying QTLs affected by local genomic differences, epistasis with the genetic background, or both.

When doing GWAS in a multi-group population, geneticists generally assume that QTL effects are conserved between groups. Such QTLs were detected in our study with the example of the SNP associated with MF in the vicinity of *Vgt2* [14] and its candidate gene: the flowering activator *ZCN8* [61–63] on chromosome 8. At this SNP, all hypotheses that tested a general SNP effect had a high −log_10_(pval), and conversely for hypotheses testing a divergent SNP effect. When simultaneously interpreting all tests, *Vgt2* appeared to have an effect that is conserved between genetic groups. Such a QTL can easily be detected in a multi-group population sample using a standard GWAS model [1]. However many QTLs show more complex patterns.

When group-specific allele effects are only due to group differences in LD or group-specific mutations at the QTL, the difference in allele effects should be conserved between the pure and the admixed genetic backgrounds. A first QTL matching this situation (*QTL4.1*) was detected by a SNP located on chromosome 4. High −log_10_(pval) were observed for the test to a divergent SNP effect between ancestries 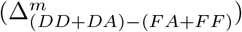, suggesting a local genomic difference. To validate this hypothesis, one could produce near isogenic lines with the two alleles from both ancestries introgressed in a dent and a flint genetic backgrounds. A phenotypic evaluation of these individuals would give a definitive proof of a local genomic difference. Nevertheless, it remains difficult to disentangle the effect of LD from that of a genetic mutation without complementary analysis. LD was shown to be different between groups, with a higher LD extent in the dent group (S4 Fig), while LD phases appeared well-conserved at short distances (S5 Fig). However, a strong overall conservation of LD phases at short distances does not exclude a specific configuration for a given SNP-QTL pair. The position of *QTL4.1* is close (< 700 Kbp) to *GRMZM2G126253*, a candidate gene for maize flowering time proposed by [59]. This gene codes for a cullin 3B protein involved in ubiquitination that was shown to be essential to plant development in *Arabidopsis* [64].

Another example is the MITE that we selected based on the *a priori* knowledge that it is associated with *Vgt1* [21, 45, 46] and its candidate gene *ZmRap2.7*. A high −log_10_(pval) was observed for a dent SNP effect (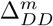 and 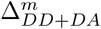) but not for a flint SNP effect. Note that another SNP (AX-91103145) was detected close to the MITE (548 Kbp further), based on 20% a FDR, for 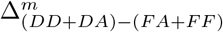 (see *QTL8.4* in S7 Fig-**a** and S4 Table). This SNP also showed evidences for a contrasted QTL effect between the dent and flint groups due to a local genomic difference. However these two loci were in very low LD with each other (below 0.05). We can reasonably suggest that the MITE and the SNP both capture a partial but different genetic information of the causal genetic variant at *Vgt1*. [46] already showed the existence of other genetic variants being more associated with maize flowering than the MITE in the vicinity of *Vgt1*, such as CGindel587.

Group-specific allele effects may also be due to an interaction with the genetic background. A first QTL matching this profile was detected by a SNP in the vicinity of *Vgt3* on chromosome 3 [53, 54] and its candidate gene *ZmMADS69* [65]. This QTL showed an effect varying according to the genetic background: large in the dent, intermediate in the admixed and small in the flint. A high −log_10_(pval) was observed for tests that supported this hypothesis: a dent SNP effect in the dent genetic background 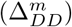 and a divergent dent SNP effect between genetic backgrounds 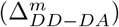. If this interaction with the background involves numerous loci, introgressing alleles from a dent to a flint genetic background may lead to disappointing results, as the effect would probably vanish with repeated back-cross generations. If interactions mostly involve a single locus, the effect at *Vgt3* effect is conditioned by the allele at the other locus, so that a simultaneous introgression may be necessary to reach the desired effect. Using near isogenic lines that cumulated an early mutation at *Vgt1* [66] and the early allele at *Vgt3*, the effect of *Vgt3* was shown to vanish in presence of the early allele of *Vgt1* (A. Charcosset pers. comm.), which supports the hypothesis of *Vgt3* interacting with the genetic background. Recently, [65] demonstrated the action of *ZmMADS69*, the candidate gene of *Vgt3*, as being an activator of the regulatory module *ZmRap2.7* - *ZCN8*, which are the candidate genes of *Vgt1* and *Vgt2*, respectively. The existence of such interactions is consistent with flowering time being controlled by a network of interacting loci, as now well established in model species arabidopis [67].

Other examples of QTLs interacting with the genetic background were identified. Two of them featured a similar profile in the sense that they mainly exhibited a QTL effect in the admixed genetic background. One was located on chromosome 2 (*QTL2.1*) and showed a flint effect in the admixed genetic background, while the other QTL was located on chromosome 7 (*QTL7.2*) and showed an opposite dent effect between the dent and the admixed genetic backgrounds. Such QTLs are interesting as they are mainly revealed when creating admixed genetic material. They also suggest complex epistatic interactions between QTLs for these traits. The position of *QTL2.1* is close (< 1.4 Mbp) to *ereb197*, a candidate gene for maize flowering time proposed by [59]. This gene codes for an AP2-EREBP transcription factor: a family of transcription factors known to play a role in plant development and response to environmental stress [68]. The position of *QTL7.2* is close (< 100 Kbp) to *dof47*, a candidate gene for maize flowering time proposed by [59]. This gene codes for a C2C2-Dof transcription factor: a family of transcription factors known to play major roles in plant growth and development [69].

The existence of epistatic interactions was also evaluated globally by a test that aimed at detecting directional epistasis [49]. This test was specifically developed to benefit from our admixed genetic material and revealed important directional epistasis for both flowering traits with admixed lines flowering closer to the dent than the flint group. Such epistasis may imply that (i) the effects of early alleles from flint origin tend to decrease in presence of alleles that are more frequent in dent than in flint group and/or (ii) the effect of late alleles from dent origin tends to be promoted by alleles that are more frequent in flint than in dent group. Alternatively, this epistasis can be interpreted as late QTL alleles (common in dent lines but rare in flint lines) interacting in a duplicate way [70], i.e. the presence of a late allele at one QTL is sufficient to confer a late phenotype. This hypothesis is equivalent to early QTL alleles (common in flint lines but rare in dent lines) interacting in a complementary way [70], i.e. early alleles are needed at both loci to confer an early phenotype. We also tested global epistasis that is not directional by decomposing the genetic variance into an additive and an epistatic component, as suggested by [71]. This confirmed the existence of epistatic interactions for FF and MF (S5 Table). In conclusion, the assessment of global epistasis supported the possibility of QTLs interacting with the genetic background, resulting from epistatic interactions with loci that have differentiated allele frequencies between groups. It would be interesting to test the existence of epistatic interactions between pairs of loci. However, a filtering on crossed allele frequencies between pairs of loci would lead to discard most SNPs from the analysis. Other possibilities would be to apply GWAS procedures that are based on testing the epistatic variance of each SNP against the polygenic background [72–74].

## Conclusion

In this study, we proposed an innovative multi-group GWAS method which accounts and tests for the heterogeneity of QTL allele effects between groups. The addition of admixed individuals to the dataset was useful to disentangle the factors causing the heterogeneity of allele effects, being either a local genomic differences or epistatic interactions with the genetic background. Only homozygous inbred lines were considered in this study, but the method may be easily generalized to heterozygous individuals. Recently many studies focused on the problem of genomic prediction across genetic groups [41, 75–78]. In such scenarios, the stability of QTL effects across genetic backgrounds is an important factor impacting the prediction accuracy. It is also an important factor of the relevancy of any marker based diagnostic in complex/structured populations. Our approach opens new perspectives to investigate this stability in a wide range of species.

## Supporting information

**S1 Fig. Imputation diagram of admixed lines.** Diagram illustrating the procedure applied to impute admixed DH lines from 15K to 600K SNPs using the parental origin of alleles.

**S2 Fig. Histogram of dent genome proportion among admixed lines.**

**S3 Fig. Genome-wide selection biases among admixed lines.** Absolute difference between observed allele frequency of the reference allele *f*_*o*_ estimated on the admixed lines and their expected value *f*_*e*_ along each chromosome (|*f*_*o*_ − *f*_*e*_|). The expected allele frequencies were computed as the mean of flint and dent allele frequencies estimated on the parental lines by taking into account the contribution of each parent. A cubic smoothing spline was adjusted using the R function “smooth.spline”, and plotted in red.

**S4 Fig. LD extent.** LD extent estimated, with a sliding window of physical distances between two pairs of loci, in dent and flint genetic groups using the average of (**a**) the standard *r*^2^ or (**b**) the 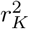 accounting for relatedness between individuals. A cubic smooth spline was adjusted for each group, using the R function “smooth.spline”.

**S5 Fig. Conservation of LD phases.** Conservation of LD phases estimated, with a sliding window of physical distances between pairs of loci, using the correlation (**a**) between the *r* of dent and flint groups (or the *r*_*K*_ accounting for relatedness between individuals), and (**b**) between the signs of r in the dent and flint groups (or the signs of the *r*_*K*_). A cubic smooth spline was adjusted for method, using the R function “smooth.spline”.

**S6 Fig. Position of QTLs detected for FF.** Position of QTLs detected for FF with a FDR of 20% using (**b**) **M**_1_, (**b**) **M**_2_ and (**c**) **M**_3_. The size of the grey dots is proportional to the −log_10_(pval) of the test at the most significant SNP of the region.

**S7 Fig. Boxplots of phenotypes adjusted for polygenic background variation using relatedness (MF K corrected)** for the different alleles of the six other highlighted QTLs: (**a**) *QTL8.4*, (**b**) *QTL10.1*, (**c**) *QTL3.5*, (**d**) *QTL6.3*, (**e**) *QTL8.6* and (**f**) *QTL2.2* using **M**_3_. The denomination of the allelic states on the x-axis include the SNP allele (0/1), its ancestry (D/F) and the genetic background in which it is observed (D/A/F), as presented in Table 2.

**S1 Table. Parameters estimated in the phenotypic analysis**. The lines “Row-Column” refer to the modeling of row and columns as defined by the experimental design. AR1 refers to the autoregressive model AR1, while IID refers to the modeling of row and column as being independent and identically distributed among rows and among columns for a given trial. For more information, see the ASReml-R reference manual by [48]. The mean of each trial *j* (with *j* ∈ {2015, 2016}) was computed following: 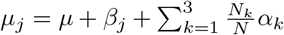 where *N*_*k*_ is the number of individuals (genotypes) in genetic background *k* (with *k* ∈ {*D*, *A*, *F*}) and *N* is the total number of individuals. The mean of each genetic background was computed following: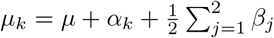. The genetic variance 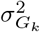 of each genetic background *k* and the GxE variance of each genetic background *k* in each trial *j* were also reported.The heritabilities of each genetic background *k* were computed as: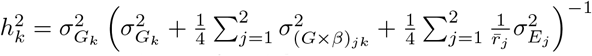 where 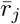 is the mean number of genotype replicates in trial *j*

**S2 Table. Information regarding significant SNPs for MF.** Information regarding significant SNPs for MF using all GWAS strategies: the name of the SNP, the chromosome on which it is located, its position in bp along the chromosome, the frequency of the allelic state observed in the dataset in which it was tested, the GWAS model applied, the hypothesis tested, the −log_10_(pval) of the test and the FDR for which it was declared significant.

**S3 Table. Information regarding significant SNPs for FF.** Information regarding significant SNPs for FF using all GWAS strategies: the name of the SNP, the chromosome on which it is located, its position in bp along the chromosome, the frequency of the allelic state observed in the dataset in which it was tested, the GWAS model applied, the hypothesis tested, the −log_10_(pval) of the test and the FDR for which it was declared significant.

**S4 Table. Information regarding the six other highlighted QTLs:** *QTL8.4*, *QTL10.1*, *QTL3.5*, *QTL6.3*, *QTL8.6* and *QTL2.2*.

**S5 Table. Additive, epistatic and residual variance components for each trait with the p-value (pval) of the epistatic component using a likelihood-ratio LR test.** The existence of epistasis can be investigated using a test based on variance components. The epistatic variance component between pairs of loci was estimated on the joint dent, flint and admixed dataset using the model: *Y*_*l*_ = *μ* + *G*_*l*_ + (*G* × *G*)_*l*_ + *E*_*l*_ where (*G* × *G*)_*l*_ is the global epistatic deviation of line *l*, all others terms being identical to those described in **M**_1_ (Eq. 1). Noting 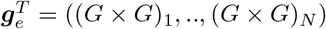, one assumes 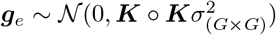 where ***K*** ○ ***K*** is the Hadamard product of the kinship matrix (Eq. 2) with itself and 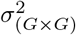 is the epistatic genetic variance between pairs of loci. This model can be seen as a simplified version of the one proposed by [71], as purely homozygous lines were used. The epistatic variance component was tested using a LR test between this model and the same model without the term (*G* × *G*)_*l*_.

## S1 Appendix. Effect of directional epistasis on the mean of an admixed progeny.

## S2 Appendix. Interpretation of the test 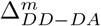.

This appendix show that 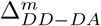 tests for an epistatic interaction between the SNP and the genetic background

## Acknowledgments

We thank Stéphane Nicolas (GQE - Le Moulon) for his contribution to genotypic data assembly. We thank Bernard Lagardère, Jean-René Loustalot (INRA Saint-Martin de Hinx) for their contribution to the panel assembly and the coordination of seed production, all the breeding companies partners of the Amaizing project for the production of admixed lines and the company Limagrain for the genotyping of admixed lines.

## Data availability statement

Genotypes, Allele ancestries, phenotypes, the R script to run analyses, and summary GWAS statistics will be available from the DRYAD (https://datadryad.org/) database (accession number, doi:). All other relevant data are within the paper and its Supporting Information files.

## Author Contributions

**Conceptualization**: Rio Simon, Mary-Huard Tristan, Moreau Laurence and Charcosset Alain

**Data Curation**: Rio Simon, Mary-Huard Tristan, Moreau Laurence, Bauland Cyril, Palaffre Carinne, Madur Delphine, Combes Valérie, Charcosset Alain

**Formal Analysis**: Rio Simon, Mary-Huard Tristan, Moreau Laurence, Charcosset Alain

**Funding Acquisition**: Charcosset Alain

**Investigation**: Rio Simon, Mary-Huard Tristan, Moreau Laurence, Bauland Cyril, Palaffre Carinne, Madur Delphine, Combes Valérie, Charcosset Alain

**Methodology**: Rio Simon, Mary-Huard Tristan, Moreau Laurence, Charcosset Alain

**Project Administration**: Charcosset Alain

**Resources**: Rio Simon, Mary-Huard Tristan, Moreau Laurence, Bauland Cyril, Palaffre Carinne, Madur Delphine, Combes Valérie, Charcosset Alain

**Software**: Rio Simon, Mary-Huard Tristan, Moreau Laurence, Charcosset Alain

**Supervision**: Charcosset Alain

**Validation**: Rio Simon, Mary-Huard Tristan, Moreau Laurence, Charcosset Alain

**Visualization**: Rio Simon, Mary-Huard Tristan, Moreau Laurence, Charcosset Alain

**Writing – Original Draft Preparation**: Rio Simon, Mary-Huard Tristan, Moreau Laurence, Charcosset Alain

