## Supplementary figures and images for "Disentangling group specific QTL allele effects from genetic background epistasis using admixed individuals in GWAS: an application to maize flowering"

### Supplementary Figure S1

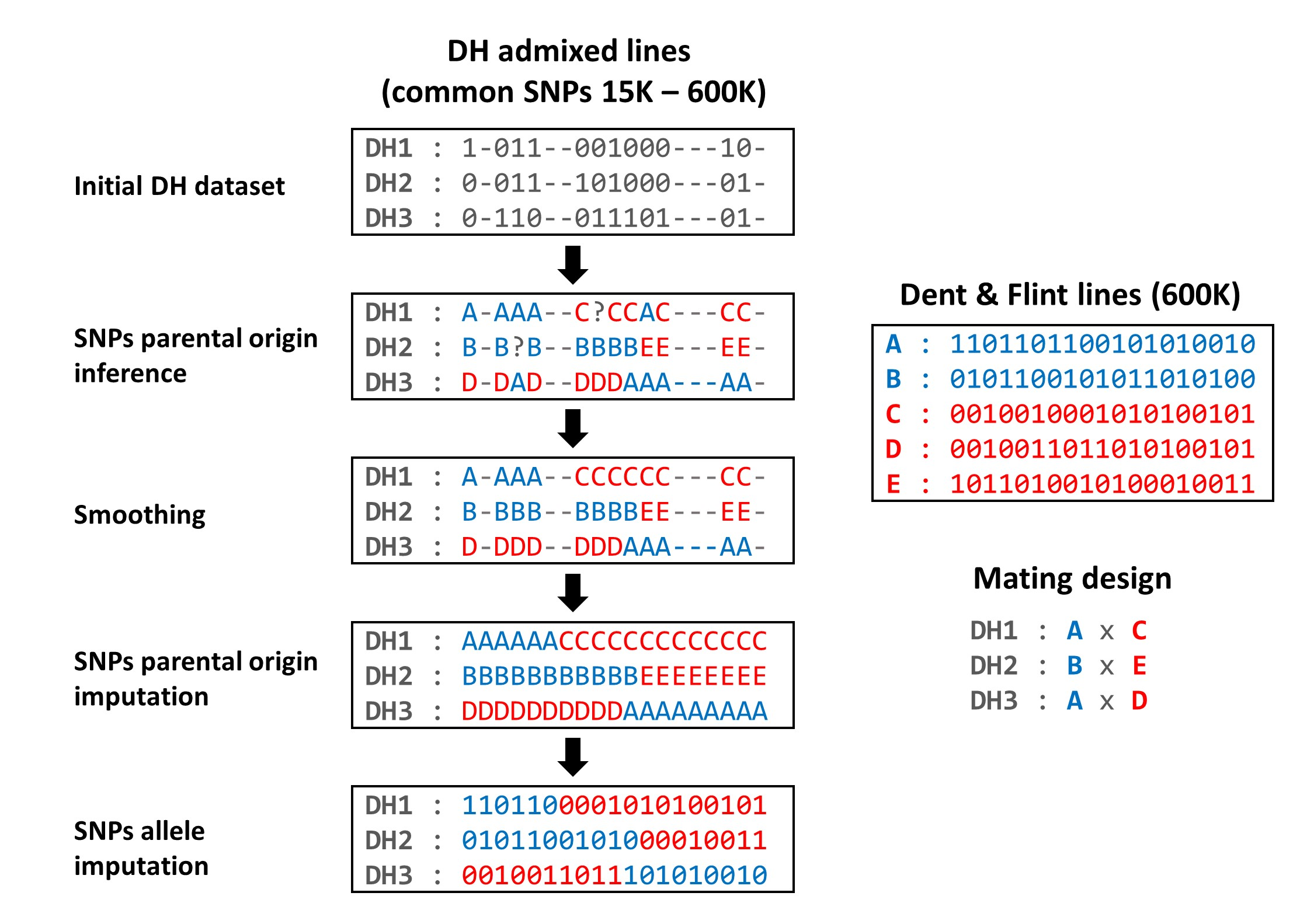

### Supplementary Figure S2

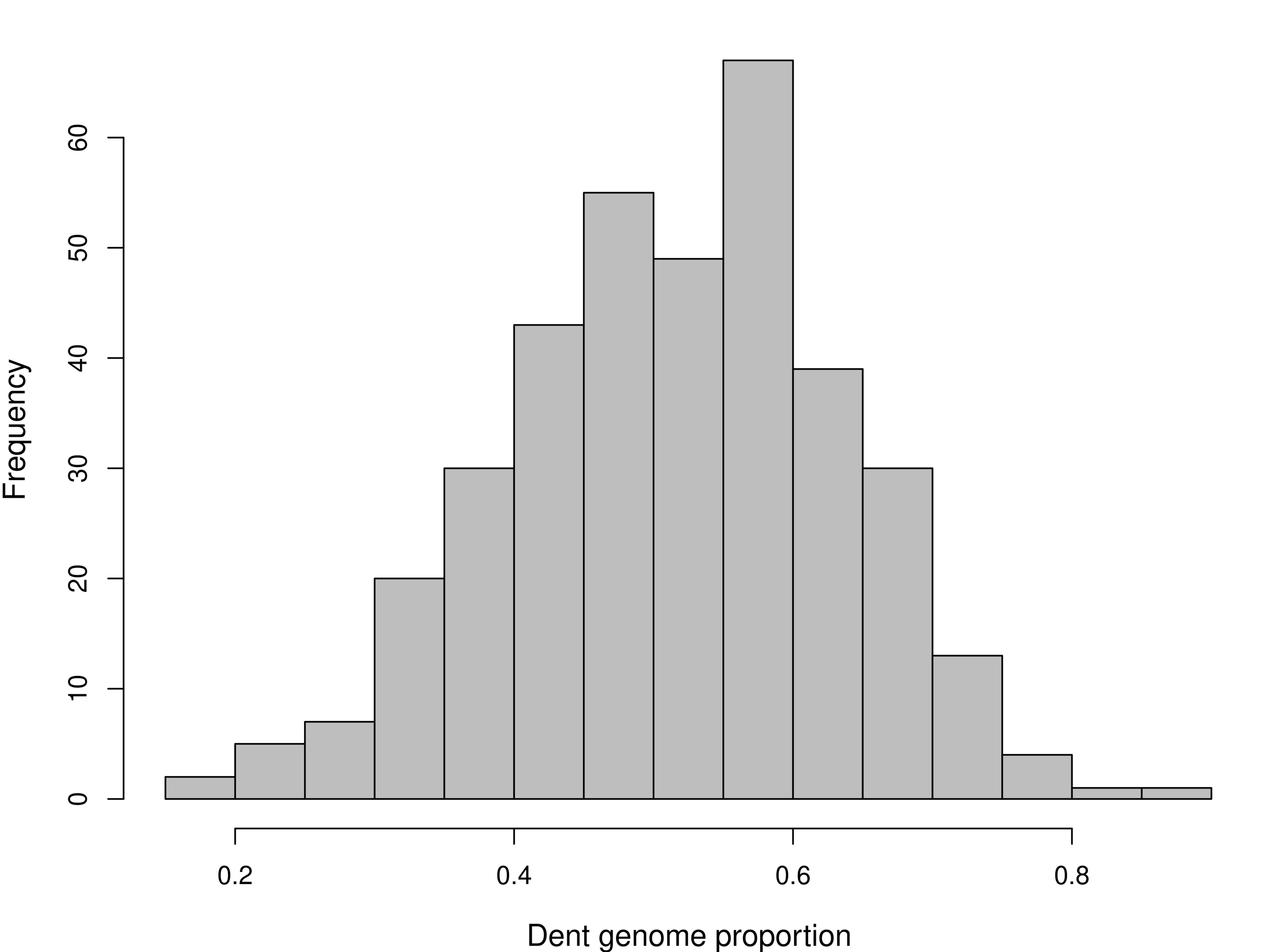

### Supplementary Figure S3

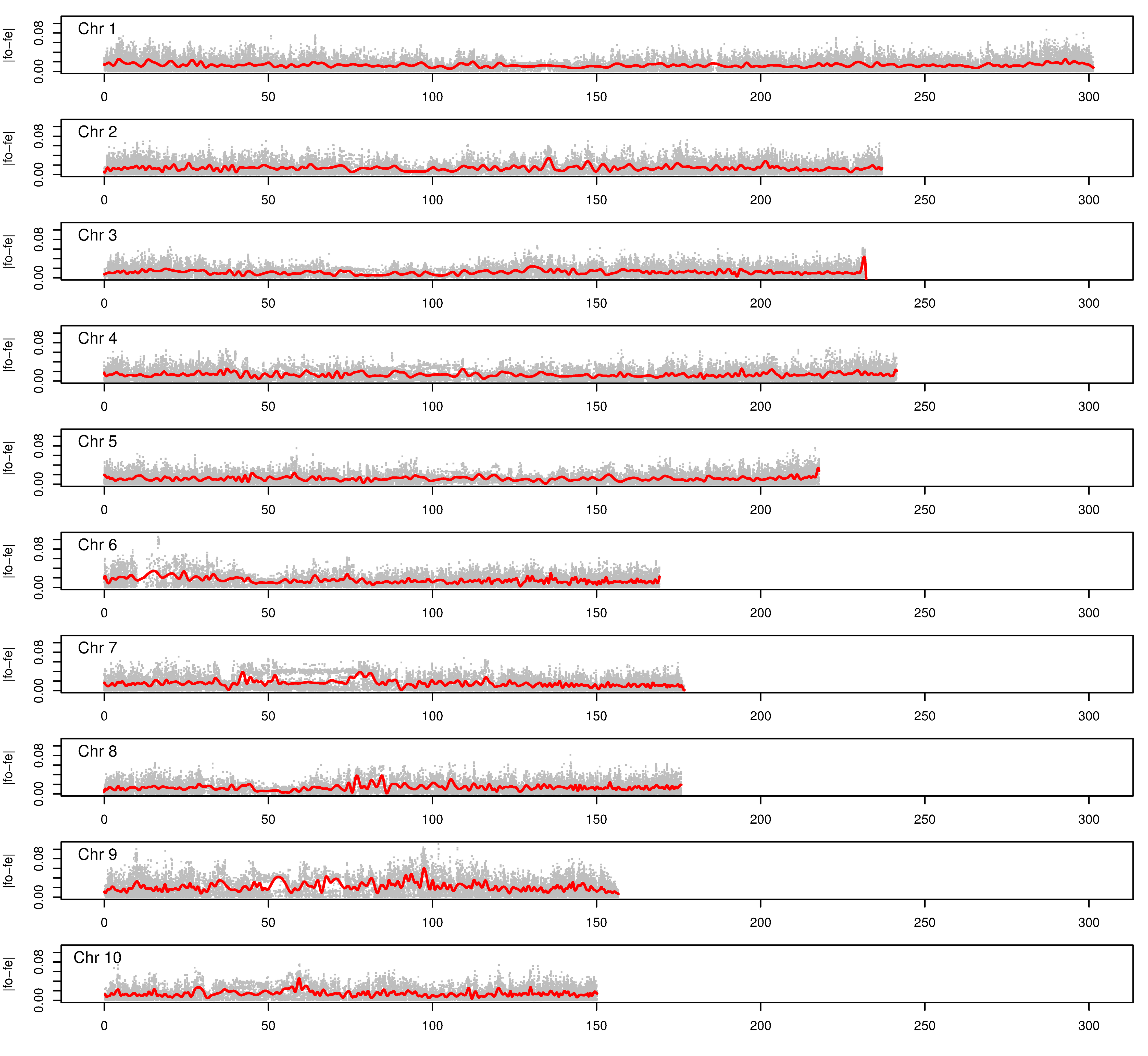

### Supplementary Figure S4

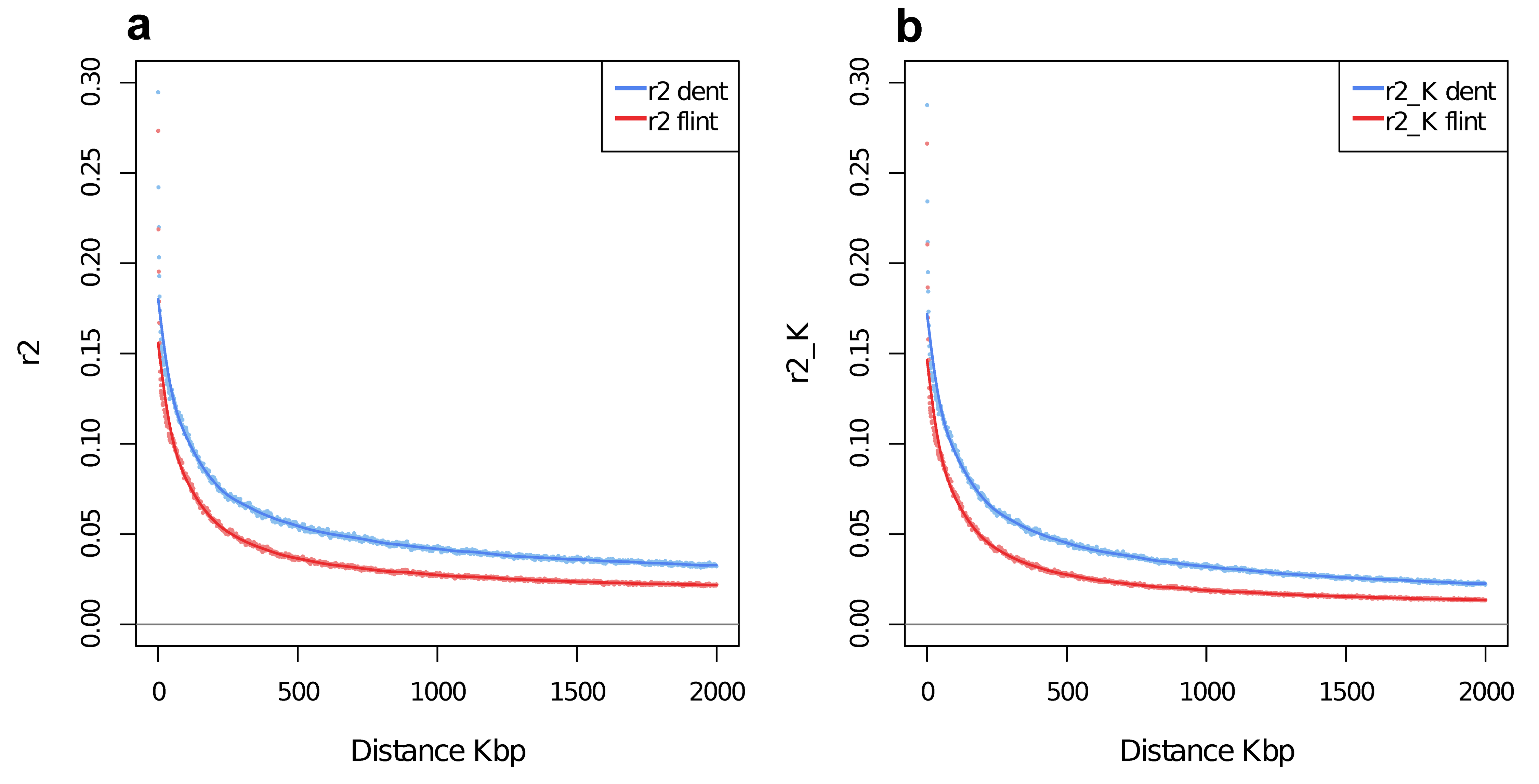

### Supplementary Figure S5

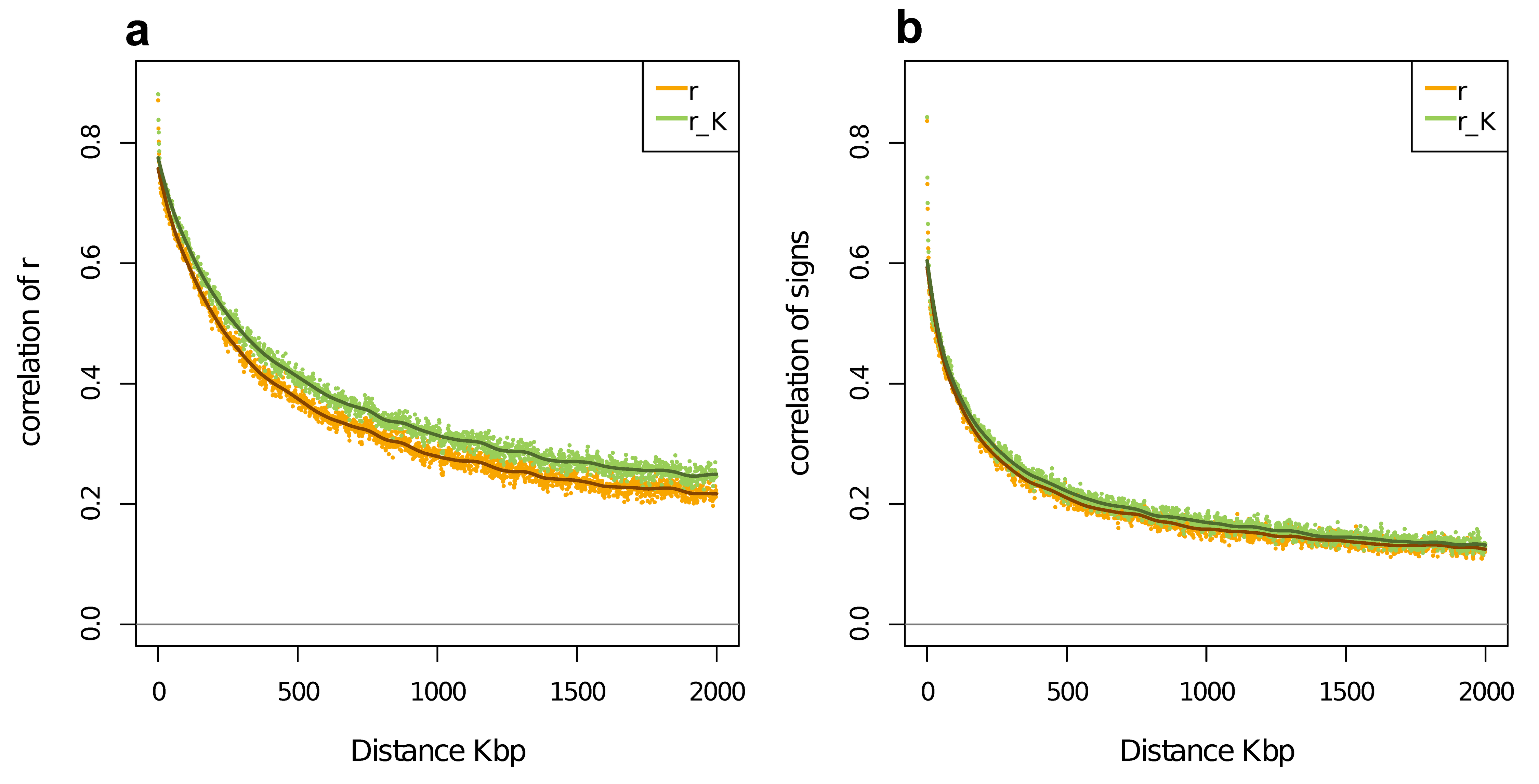

### Supplementary Figure S6

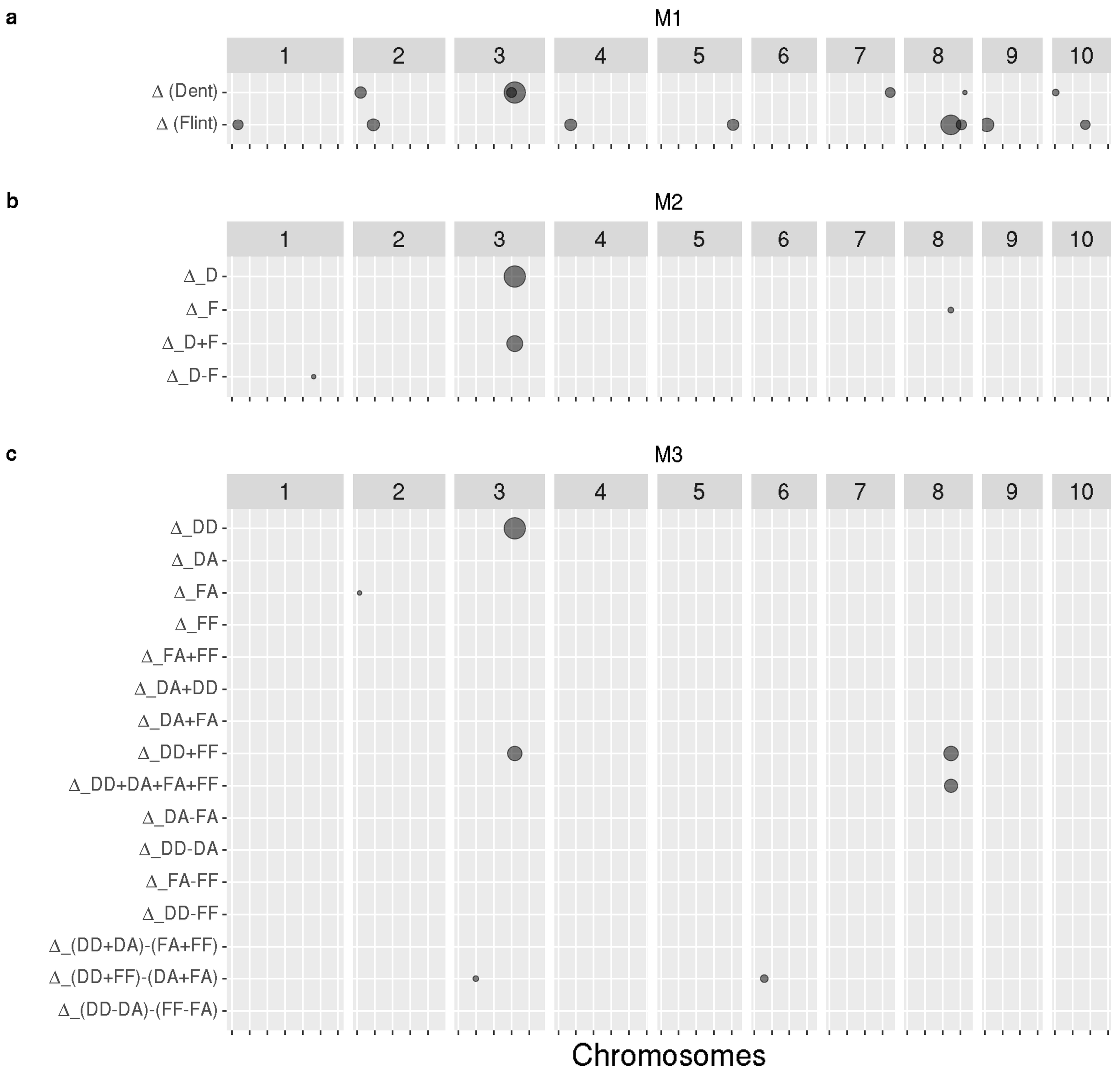

### Supplementary Figure S7

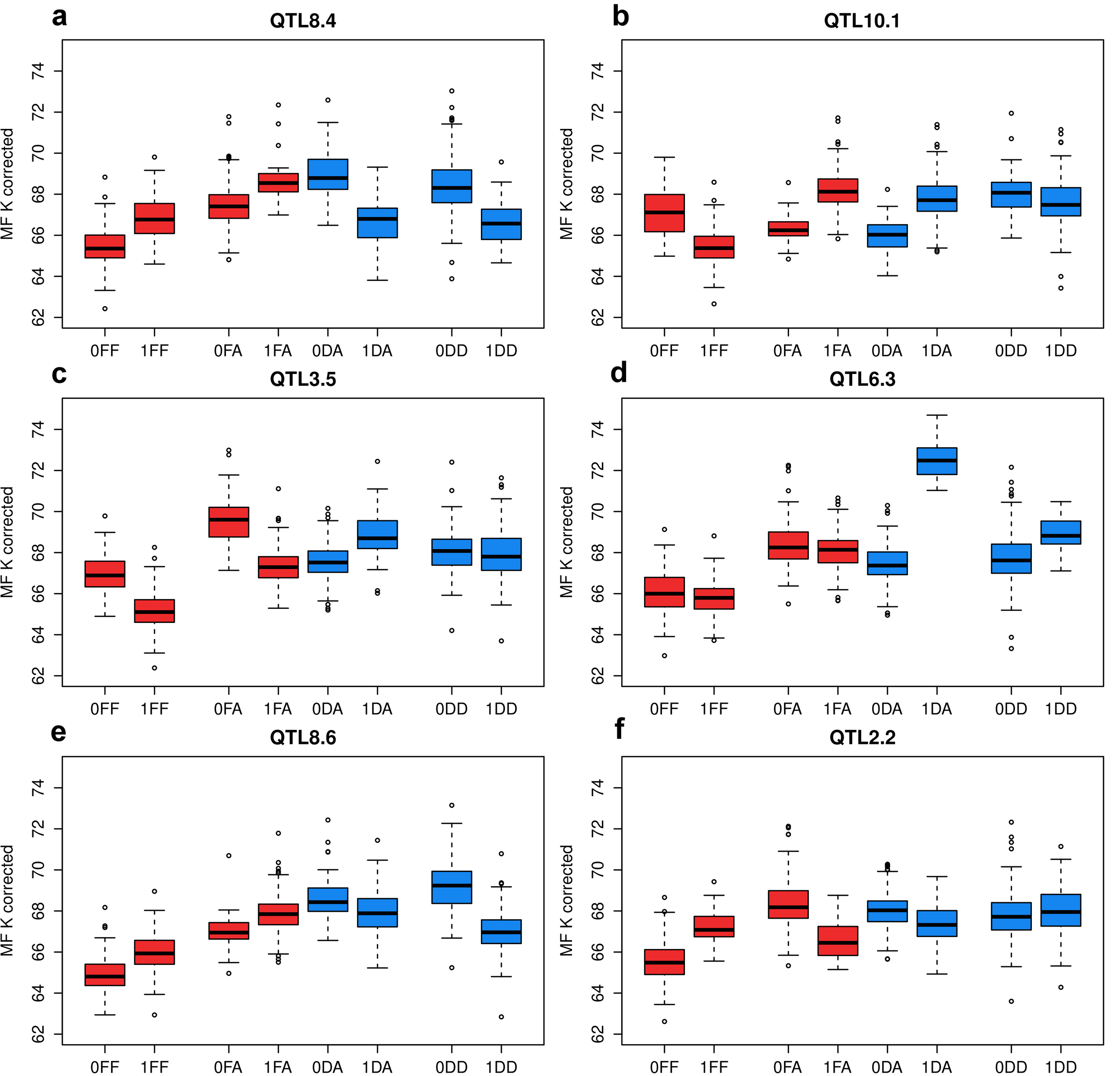
