## Appendix S1 for "Disentangling group specific QTL allele effects from genetic background epistasis using admixed individuals in GWAS: an application to maize flowering"

### S1 Appendix

#### Effect of directional epistasis on the mean of an admixed progeny

Genetic values are modeled using allele effects at bi-allelic QTLs and deviation effects for each pair of alleles coming from different loci, leading to epistatic interactions between QTLs:

$$G_i = \sum_{m=1}^{M} [(1 - W_{im})\beta_m^0 + W_{im}\beta_m^1] + \sum_{m=1}^{M} \sum_{m'>m}^{M} [(1 - W_{im})(1 - W_{im'})\delta_{mm'}^{00} + W_{im}(1 - W_{im'})\delta_{mm'}^{10} + (1 - W_{im})W_{im'}\delta_{mm'}^{01} + W_{im}W_{im'}\delta_{mm'}^{11}]$$

where $G_i$ is the genotype of individual $i$, $M$ is the number of QTLs, $W_{im}$ is the QTL genotypes of individual $i$ at locus $m$ (coded 0/1) with $W_{im} \sim \mathcal{B}(f_m)$ independent, $f_m$ is allele frequency of allele 1, $\beta_m^0$ is effect of allele 0 at locus $m$, $\beta_m^1$ is effect of allele 1 at locus $m$, $\delta_{mm'}^{00}$ is the deviation effect specific to the pair of alleles 0 at locus $m$ and $m'$, $\delta_{mm'}^{10}$ is the deviation effect specific to the pair of allele 0 at locus $m$ and allele 1 at locus $m'$, $\delta_{mm'}^{01}$ is the deviation effect specific to the pair of allele 1 at locus $m$ and allele 1 at locus $m'$ and $\delta_{mm'}^{11}$ is the deviation effect specific to the pair of alleles 1 at locus $m$ and $m'$.

Let $\mu$ be the expected value of $G_i$:

$$\mu = \sum_{m=1}^{M} [\beta_m^0 + f_m(\beta_m^1 - \beta_m^0)] + \sum_{m=1}^{M} \sum_{m'>m}^{M} [\delta_{mm'}^{00} + f_m(\delta_{mm'}^{10} - \delta_{mm'}^{00}) + f_{m'}(\delta_{mm'}^{01} - \delta_{mm'}^{00}) + f_m f_{m'}(\delta_{mm'}^{11} + \delta_{mm'}^{00} - \delta_{mm'}^{10} - \delta_{mm'}^{01})]$$

Let us consider two genetic groups D and F of pure lines and a group of admixed lines A. Both group D and F have specific allele frequencies $f_{mD}$ and $f_{mF}$ at each locus $m$. Let us suppose that admixed lines were obtained from across-group hybrids without selection ($f_{mA} = \frac{f_{mD} + f_{mF}}{2}$ at a given locus $m$). Then, the mean difference between $\mu_A$ and the expected mean $\frac{\mu_D + \mu_F}{2}$ is:

$$\mu_A - \frac{\mu_D + \mu_F}{2} = -\frac{1}{4} \sum_{m=1}^{M} \sum_{m'>m}^{M} (f_{mD} - f_{mF})(f_{m'D} - f_{m'F}) \left(\delta_{mm'}^{11} + \delta_{mm'}^{00} - \delta_{mm'}^{10} - \delta_{mm'}^{01}\right)$$

In absence of epistatic interactions between loci, this mean difference is null. In presence of epistatic interactions between loci, the mean difference is not null if epistasis is directional, i.e. if QTL deviation effects do not cancel each other out among loci. Moreover, the higher the differentiation in allele frequencies between groups D and F for a given pair of loci, the higher the contribution of this pair to the mean difference.
