## Appendix S2 for "Disentangling group specific QTL allele effects from genetic background epistasis using admixed individuals in GWAS: an application to maize flowering"

### S2 Appendix

#### Interpretation of the test $\Delta^m_{DD-DA}$

Let us re-parametrize model $\mathbf{M_3}$ so that

$$\beta^m_{ijk} = \alpha_i + \gamma_j + \delta_k + (\alpha\gamma)_{ij} + (\alpha\delta)_{ik} + (\gamma\delta)_{jk} + (\alpha\gamma\delta)_{ijk}$$

where:

- $\alpha_i$ : effect of genotypic allele $i$ with $i \in \{0, 1\}$
- $\gamma_j$ : effect of allele ancestry $j$ with $j \in \{D, F\}$
- $\delta_k$ : effect of genetic background $k$ with $k \in \{D, A, F\}$
- $(\alpha\gamma)_{ij}$ : effect of the "genotypic allele - allele ancestry" interaction
- $(\alpha\delta)_{ik}$ : effect of the "genotypic allele - genetic background" interaction
- $(\gamma\delta)_{jk}$ : effect of the "allele ancestry - genetic background" interaction
- $(\alpha\gamma\delta)_{ijk}$ : effect of the "genotypic allele - allele ancestry - genetic background" interaction

Let us write the test $\Delta^m_{DD-DA}$ according to the new parametrization:

$$\begin{aligned}
\Delta^m_{DD-DA} &= (\beta^m_{1DD} - \beta^m_{0DD}) - (\beta^m_{1DA} - \beta^m_{0DA}) \\
\Delta^m_{DD-DA} &= (\alpha_1 + \gamma_D + \delta_D + (\alpha\gamma)_{1D} + (\alpha\delta)_{1D} + (\gamma\delta)_{DD} + (\alpha\gamma\delta)_{1DD} \\
&\quad -\alpha_0 - \gamma_D - \delta_D - (\alpha\gamma)_{0D} - (\alpha\delta)_{0D} - (\gamma\delta)_{DD} - (\alpha\gamma\delta)_{0DD}) \\
&\quad - (\alpha_1 + \gamma_D + \delta_A + (\alpha\gamma)_{1D} + (\alpha\delta)_{1A} + (\gamma\delta)_{DA} + (\alpha\gamma\delta)_{1DD} \\
&\quad -\alpha_0 - \gamma_D - \delta_A - (\alpha\gamma)_{0D} - (\alpha\delta)_{0A} - (\gamma\delta)_{DA} - (\alpha\gamma\delta)_{0DA}) \\
\Delta^m_{DD-DA} &= ((\alpha\delta)_{1D} - (\alpha\delta)_{0D}) - ((\alpha\delta)_{1A} - (\alpha\delta)_{0A}) \\
&\quad + ((\alpha\gamma\delta)_{1DD} - (\alpha\gamma\delta)_{0DD}) - ((\alpha\gamma\delta)_{1DA} - (\alpha\gamma\delta)_{0DA})
\end{aligned}$$

In this last expression, all terms involve an interaction between the genotypic allele and the genotypic background.
